# Comprehensive mapping of mutations to the SARS-CoV-2 receptor-binding domain that affect recognition by polyclonal human serum antibodies

**DOI:** 10.1101/2020.12.31.425021

**Authors:** Allison J. Greaney, Andrea N. Loes, Katharine H.D. Crawford, Tyler N. Starr, Keara D. Malone, Helen Y. Chu, Jesse D. Bloom

## Abstract

The evolution of SARS-CoV-2 could impair recognition of the virus by human antibody-mediated immunity. To facilitate prospective surveillance for such evolution, we map how convalescent serum antibodies are impacted by all mutations to the spike’s receptor-binding domain (RBD), the main target of serum neutralizing activity. Binding by polyclonal serum antibodies is affected by mutations in three main epitopes in the RBD, but there is substantial variation in the impact of mutations both among individuals and within the same individual over time. Despite this inter- and intra-person heterogeneity, the mutations that most reduce antibody binding usually occur at just a few sites in the RBD’s receptor binding motif. The most important site is E484, where neutralization by some sera is reduced >10-fold by several mutations, including one in emerging viral lineages in South Africa and Brazil. Going forward, these serum escape maps can inform surveillance of SARS-CoV-2 evolution.

## Introduction

Neutralizing antibodies against the SARS-CoV-2 spike are associated with protection against infection in both humans (Addetia et al., 2020; Lumley et al., 2020) and animal models (Alsoussi et al., 2020; Walls et al., 2020; Zost et al., 2020a). However, other human coronaviruses undergo antigenic evolution that erodes neutralizing antibody immunity (Eguia et al., 2020). This antigenic evolution is driven by positive selection for mutations in the viral spike, particularly in regions involved in receptor binding (Kistler and Bedford, 2020; Wong et al., 2017). To monitor for similar antigenic evolution of SARS-CoV-2, it is important to determine which viral mutations impact human polyclonal antibody immunity.

A multitude of recent studies have identified viral mutations that escape monoclonal antibodies targeting the SARS-CoV-2 spike (Baum et al., 2020; Greaney et al., 2020; Li et al., 2020; Liu et al., 2020b; Starr et al., 2020a; Weisblum et al., 2020). However, it remains unclear how mutations that escape specific monoclonal antibodies will affect the polyclonal antibody response elicited by infection or vaccination. Several recent studies have identified viral mutations that impact neutralization by polyclonal human sera. So far, these studies have relied on either selecting viral escape mutants with reduced neutralization sensitivity (Andreano et al., 2020; Weisblum et al., 2020), or characterizing the antigenic effects of specific mutations such as those observed in circulating viral isolates (Kemp et al., 2020b; Li et al., 2020; Liu et al., 2020b; Thomson et al., 2020). This work has shown that single mutations to the spike’s receptor-binding domain (RBD) or N-terminal domain (NTD) can appreciably reduce viral neutralization by polyclonal sera, sometimes by as much as 10-fold. However, a limitation of these studies is that they characterize an incomplete subset of all possible mutations, and thus do not completely describe the effects of viral mutations on recognition by polyclonal serum antibodies.

Here we comprehensively map how all amino-acid mutations to the SARS-CoV-2 spike RBD affect binding by the antibodies in plasma collected from convalescent individuals ~1 to ~3 months post-symptom onset. We focus on the RBD because prior studies have reported that RBD-binding antibodies contribute the majority of the neutralizing activity of most human sera (Piccoli et al., 2020; Steffen et al., 2020), a result we confirm. Our complete maps of how mutations impact serum antibody binding identify three major epitopes in the RBD. However, serum antibody binding from different individuals is impacted differently by mutations in these epitopes, and sometimes the impacts of mutations also change over time for longitudinal samples from the same individual. Some mutations that reduce serum antibody binding also reduce viral neutralization by >10 fold. The site where mutations tend to have the largest effect on binding and neutralization is E484, which unfortunately is a site where mutations are present in several emerging SARS-CoV-2 lineages (Tegally et al., 2020; Voloch et al., 2020). However, some sera are more affected by mutations at other sites, while others are largely unaffected by any single mutation. Overall, these systematic maps of how mutations to the SARS-CoV-2 RBD affect recognition by human antibody immunity can inform surveillance of ongoing viral evolution.

## Results

### RBD-targeting antibodies dominate the neutralizing activity of most convalescent sera

We characterized the serum antibodies from 35 plasma samples longitudinally collected from 17 different SARS-CoV-2-infected individuals between 15 and 121 days post-symptom onset. Prior work has shown that these samples all have RBD-binding antibodies and neutralizing activity, with a median neutralization titer 50% (NT50) of ~250 (range of 34 to >10,000) against lentiviral particles pseudotyped with the D614 variant of the SARS-CoV-2 spike. For most of the 17 individuals, both the RBD binding and the neutralizing activity decreased moderately from one to four months post-infection (Crawford et al., 2020a) (**Supplementary Table 1**).

Several recent studies have reported that RBD-binding antibodies contribute the majority of the neutralizing activity in most convalescent human sera (Piccoli et al., 2020; Steffen et al., 2020). To confirm the importance of anti-RBD antibodies for the samples in our study, we used RBD-conjugated beads to deplete RBD-binding antibodies, and compared the neutralizing activity pre- and post-depletion. First, we validated that these depletions effectively removed RBD-directed but not other anti-spike antibodies. To do this, we created “synthetic” sera by combining non-neutralizing pre-pandemic serum with either an RBD-binding or N-terminal domain (NTD)-binding monoclonal antibody. As expected, RBD antibody depletion completely eliminated binding of the anti-RBD synthetic serum to both RBD and spike, but did not reduce the spike binding activity of the anti-NTD synthetic serum (**Figure S1A**).

We then validated that the depletion effectively removed nearly all RBD-binding antibodies from the convalescent human plasma samples, and examined how depletion of RBD-binding antibodies affected total serum binding to the spike ectodomain (**Figures 1A, S1B**). To do this, we performed ELISAs for binding to the RBD and spike for each sample both pre- and post-depletion, using an anti-IgG secondary antibody. The depletion removed essentially all RBD-binding IgG antibodies but only modestly decreased the amount of IgG that bound to spike (**Figures 1A, S1B**). This result suggests that RBD-binding antibodies comprise a relatively modest proportion of all spike-binding IgG serum antibodies in naturally infected individuals, consistent with studies reporting that less than half of spike-reactive B cells and monoclonal antibodies bind to RBD (Brouwer et al., 2020; Huang et al., 2020; Seydoux et al., 2020; Voss et al., 2020).

**Figure 1.**
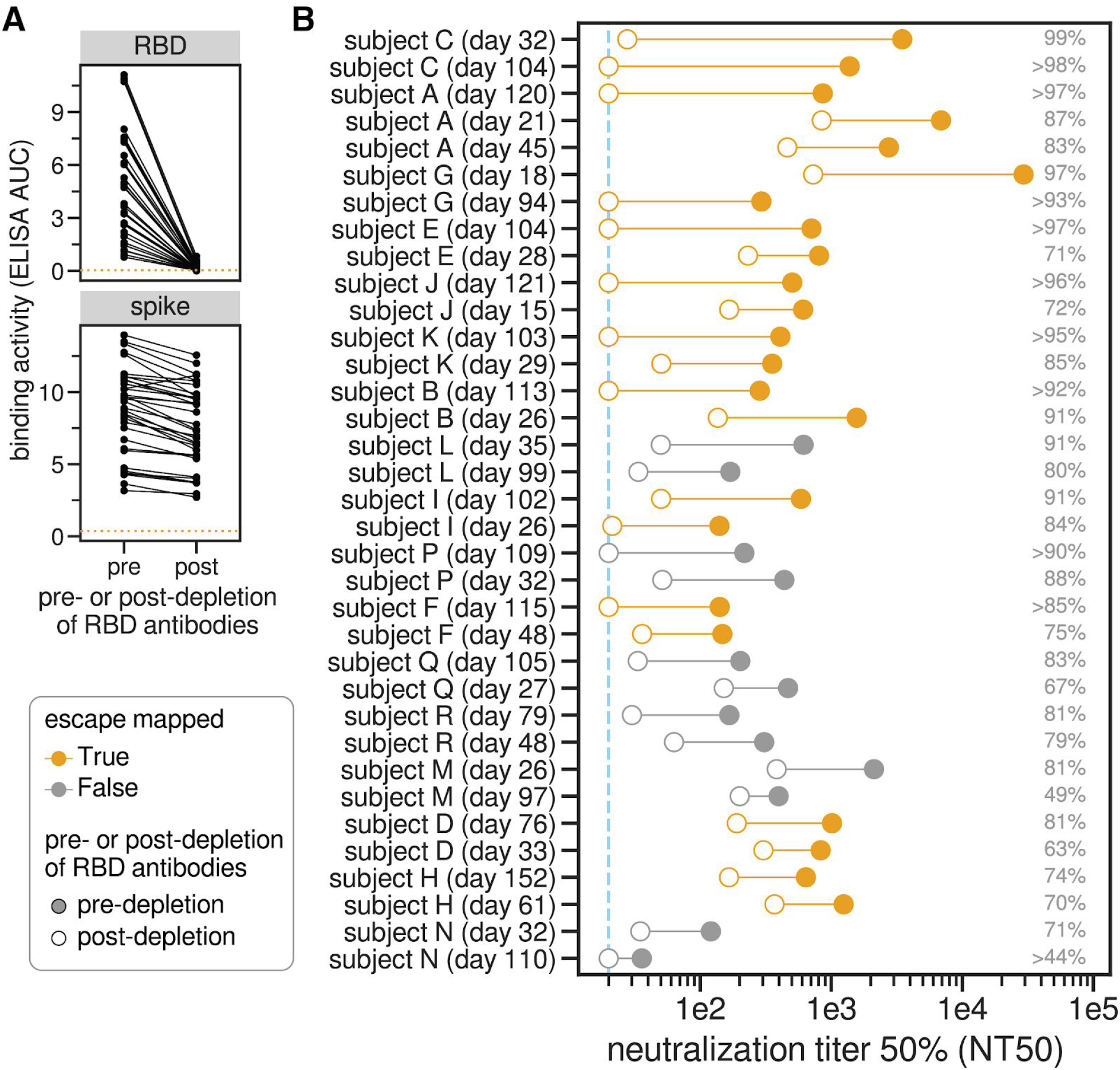
RBD-binding antibodies are responsible for most of the neutralizing activity of human polyclonal sera. **(A)** Change in binding of human sera to RBD and spike before and after depletion of RBD antibodies, measured by ELISA area under the curve (AUC). The dashed orange line is binding of pre-pandemic pooled sera collected in 2017-2018. Raw ELISA binding curves in **Figure S1A. (B)** Neutralization titer 50% (NT50) of human sera before and after depletion of RBD-binding antibodies. Legend is at left: filled and open circles are pre- and post-depletion samples, respectively, connected by a line. Orange indicates sera for which we subsequently mapped mutations that reduce binding. The numbers at right indicate the percent of all neutralizing activity attributable to RBD-binding antibodies. Sera are sorted in descending order of percent of neutralization due to RBD-binding antibodies, first by subject and then within subject. The dashed blue line is the limit of detection (NT50 of 20). Points on this line have an NT50 <= 20, so the percent of neutralization due to RBD-binding antibodies may be an underestimate for these sera. See **Figure S1** and **Supplementary Table 1** for additional data including full ELISA and neutralization curves and numerical values plotted here.

We next measured how depletion of RBD-binding antibodies affected neutralization of lentiviral particles pseudotyped with the G614 variant of the SARS-CoV-2 spike, and found that RBD-binding antibodies usually dominated the neutralizing activity (**Figures 1B, S1C, Supplementary Table 1**). Specifically, the majority of the neutralizing activity was due to RBD-binding antibodies in nearly all samples (33 of 35 tested), and >90% of neutralizing activity was due to RBD-binding antibodies in over a third of the samples (13 of 35 tested) (**Figures 1B, S1C**).

Notably, RBD-binding antibodies dominated the serum neutralizing activity both at early (~30 day) and late (~100 day) time points post-symptom onset. For many individuals, the contribution of RBD-binding antibodies to serum neutralizing activity increased over time, although this was not always the case (**Figures 1B, S1E**). For instance, the contribution of RBD-binding antibodies to neutralization increased over time for subjects E and J, but not subjects L or M. The strong contribution of RBD-binding antibodies to serum neutralization demonstrates that mapping mutations that escape these antibodies is crucial for understanding the potential for SARS-CoV-2 antigenic evolution.

### Complete mapping of RBD mutations that reduce binding by sera collected ~1 month post-symptom onset

To completely map RBD mutations that reduce binding by polyclonal serum antibodies, we extended a deep-mutational scanning method previously developed to identify mutations that escape binding by monoclonal antibodies (Greaney et al., 2020). Briefly, we used libraries of yeast that each expressed a different RBD variant on their surface. The library covered nearly all possible single amino-acid mutations to the RBD (Starr et al., 2020b). We incubated these yeast libraries with polyclonal human sera, and used fluorescence-activated cell sorting (FACS) with an IgG/IgA/IgM secondary antibody to enrich for yeast expressing RBD mutants that bound appreciably less serum antibodies than unmutagenized RBD (**Figure S2A-C**). We then used deep sequencing to measure the frequency of each RBD mutation in the initial population and the antibody-escape FACS bin. We quantified the effect of each RBD mutation on serum antibody binding as that mutation’s “escape fraction,” which is the fraction of all yeast cells expressing RBD with that mutation that fall into the FACS escape bin. These escape fractions range from 0 (no effect on serum antibody binding) to 1 (all cells with this mutation are in the antibody-escape bin) (**Figure S2A-C**) (Greaney et al., 2020; Starr et al., 2020b). All mapping experiments were performed in biological duplicate using independently constructed RBD mutant libraries; the replicates were highly correlated (**Figure S2D**,**E**), and we report the average measurements across the two libraries throughout.

We began by mapping mutations that reduced binding by serum antibodies in samples collected from 11 individuals at approximately 30 days post-symptom onset (range 15–61 days). These samples had neutralizing titers against lentiviral particles pseudotyped with the G614 variant of the SARS-CoV-2 spike that ranged from 140 to 30,000, with the extent of neutralization attributable to RBD-targeting antibodies ranging from 63% to 99% (first time point for subjects shown in orange in **Figure 1B**). We quantified each RBD mutation’s escape fraction and visualized the results using “escape maps,” which are logo plots where the height of each letter is proportional to that mutation’s escape fraction (**Figure 2A, S2A**). Interactive versions of these escape maps are at https://jbloomlab.github.io/SARS-CoV-2-RBD_MAP_HAARVI_sera.

**Figure 2.**
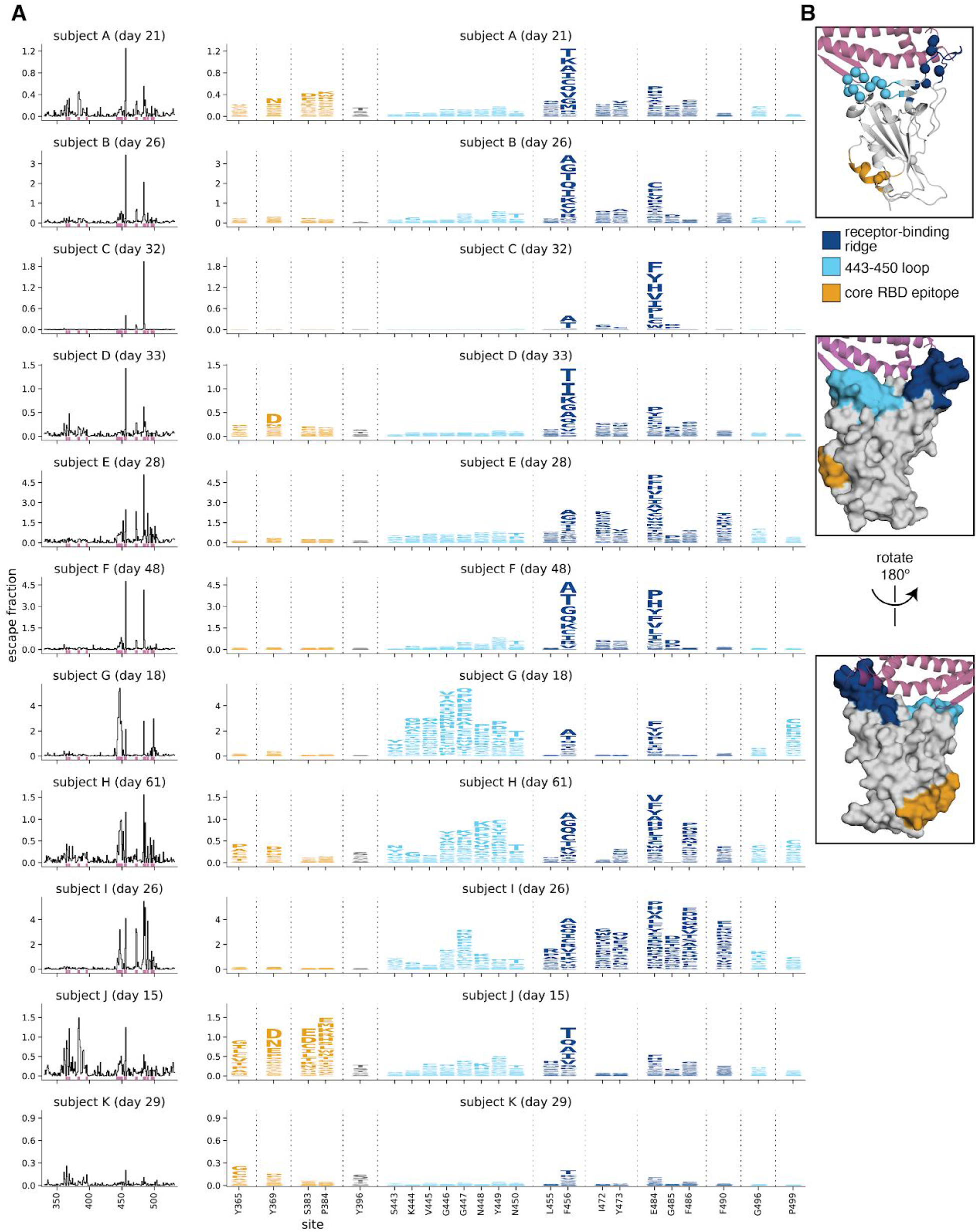
Complete maps of RBD mutations that reduce binding by polyclonal serum antibodies from 11 individuals. **(A)** The line plots at left indicate the total effect of all mutations at each site in the RBD on serum antibody binding, with larger values indicating a greater reduction in antibody binding. The logo plots at right zoom in on individual mutations at key sites (indicated by purple highlighting on the x-axis of the line plots). In these logo plots, the height of each letter is that mutation’s escape fraction, so larger letters indicate mutations that cause a greater reduction in antibody binding. Escape fractions are comparable across sites within a sample, but not necessarily between samples due to the use of sample-specific FACS gates—therefore, for each serum, the y-axis is scaled independently (see **Methods**). Sites in the logo plots are colored by RBD epitope. **(B)** For coloring of the logo plots, we designated three RBD epitopes based on the structural locations where mutations had large effects on serum antibody binding. The images show the structure of the RBD bound to ACE2 (PDB 6M0J, (Lan et al., 2020)) in several representations. The receptor-binding-ridge epitope is dark blue, the epitope containing the 443–450 loop is cyan, the core-RBD epitope is orange, the rest of the RBD is gray, and ACE2 is purple. For the cartoon rendering in the top structure, alpha carbons for sites of strong binding-escape for any of the 11 sera (i.e., all sites shown in the logo plots) are represented as spheres. Interactive versions of these escape maps are available at https://jbloomlab.github.io/SARS-CoV-2-RBD_MAP_HAARVI_sera/.

Although the exact effects of mutations on serum antibody binding varied widely across individuals, the escape maps revealed several common patterns. Mutations that strongly reduced binding fell in one of three discrete regions of the RBD: the receptor-binding ridge within the receptor-binding motif (RBM), a loop in the RBM opposite the ridge (spanning sites 443–450, and the structurally adjacent sites at 494–501), or a surface patch in the core RBD (**Figure 2B**). The receptor-binding ridge and 443–450 loop are also targeted by many potently neutralizing antibodies, including the two antibodies in the Regeneron cocktail (Greaney et al., 2020; Hansen et al., 2020; Starr et al., 2020a). The core RBD epitope is targeted by monoclonal antibodies that tend to be less potently neutralizing but more broadly cross-reactive to SARS-like coronaviruses (Barnes et al., 2020a; Piccoli et al., 2020; Yuan et al.; Zost et al., 2020a). In particular, binding by all 11 samples was reduced by mutations at site F456, and binding by most samples (9 of 11) was reduced by mutations at site E484 (**Figure 2A**). Both of these sites are within the receptor-binding ridge epitope. Notably, E484 is a site at which mutations have recently been demonstrated to reduce neutralization by several monoclonal antibodies and sera (Andreano et al., 2020; Greaney et al., 2020; Starr et al., 2020a; Weisblum et al., 2020).

We grouped the samples into several classes based on which mutations most strongly reduced serum antibody binding (**Figure 3** and the interactive visualizations at https://jbloomlab.github.io/SARS-CoV-2-RBD_MAP_HAARVI_sera/). Binding by 6 of the 11 samples was reduced primarily by mutations in the receptor-binding ridge. For instance, binding by serum antibodies from subject B (day 26) was most strongly affected by mutations at sites F456 and E484 (**Figure 2, 3A**). Binding by three samples was strongly reduced by mutations across a broader swath of the RBM, including the 443–450 loop (**Figure 3B**). An example is subject G (day 18), which was strongly affected by mutations at sites 443–450 in addition to F456 and E484. Binding by two samples was most affected by mutations in the core RBD epitope (**Figure 3C**). The sites where mutations reduced binding by these core-RBD targeting sera clustered around the lipid-binding pocket in the RBD, where binding of free fatty acids may contribute to locking spike into a “closed” conformation (Carrique et al., 2020; Toelzer et al., 2020). Notably, for the sample from subject K (day 29), no single RBD mutation had more than a small effect on serum antibody binding (**Figures 2, 3D**).

**Figure 3.**
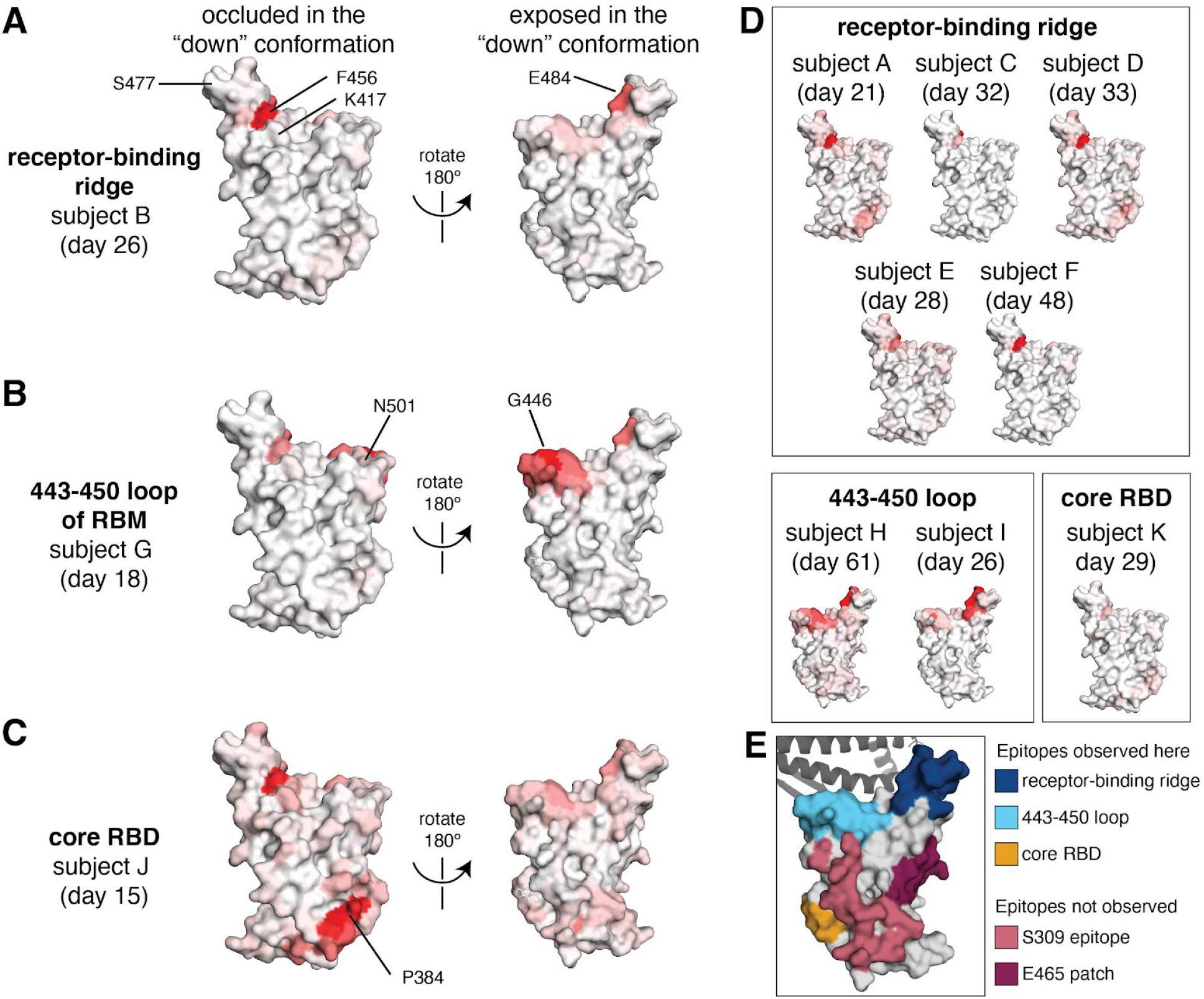
Regions of the RBD where mutations strongly reduced binding by the antibodies in sera collected from 11 individuals. The total effect of mutations at each site (sum of escape fractions) are projected onto the structure of the RBD (PDB 6M0J), with white indicating no effect of mutations at that site and red indicating a large reduction in antibody binding. Two views of the RBD are shown: the surface of the RBD that is buried in the “down” conformation, and the surface that is always exposed and accessible (Walls et al., 2020; Wrapp et al., 2020). **(A)** For some individuals (typified by subject B), antibody binding is predominantly reduced by mutations in the receptor-binding ridge, particularly at sites F456 and E484. **(B)** For some individuals (typified by subject G), antibody binding is strongly reduced by mutations in the 443–450 loop of the RBM in addition to the receptor-binding ridge. **(C)** For a few individuals (typified by subject J), antibody binding is affected by mutations in the core RBD epitope around site P384. **(D)** Samples from the other eight individuals fall in one of the three classes detailed in panels **(A)** to **(C)**. For panels **(A)** to **(D)**, the white-to-red coloring scale is set to span the same range as the y-axis limits for that serum in **Figure 2. (E)** Mutations in two major surface regions (the S309 epitope and the sites near E465) do not strongly affect serum antibody binding for any of the subjects. Shown is a surface representation of the RBD, with the 3 polyclonal serum epitopes colored as in **Figure 2**. The S309 epitope and region near E465 (“E465 patch”) are shown in pink and maroon. ACE2 is shown in a dark gray cartoon representation. Interactive versions of these structural visualizations are available at https://jbloomlab.github.io/SARS-CoV-2-RBD_MAP_HAARVI_sera/.

There are some regions of the RBD where mutations did not strongly affect serum antibody binding for any sample in our panel. These regions include the sites near the 343 glycan that are targeted by the SARS-CoV-1 cross-reactive antibody S309 (Pinto et al., 2020), and the region near residue E465 on the “lateral” side of the RBD, which to our knowledge is not an epitope for any known neutralizing antibodies (**Figure 3E**) (Barnes et al., 2020a; Greaney et al., 2020; Piccoli et al., 2020; Starr et al., 2020b). Antibodies targeting these two regions may be rare, have low binding avidity in the context of polyclonal serum, or be subdominant relative to other RBD epitopes (Barnes et al., 2020b; Piccoli et al., 2020; Weisblum et al., 2020).

### How mutations affect serum antibody binding can shift over time in the same individual

Next, we examined how the RBD mutations that affect serum antibody binding change over time as an individual’s immune response matures. We speculated that such changes might occur because other studies have shown that anti-SARS-CoV-2 antibodies become more somatically hypermutated and less clonal in the months following recovery from infection (Gaebler et al., 2020; Nielsen et al., 2020; Rodda et al., 2020). Moreover, we reasoned that mapping mutations that affect serum antibody binding several months after infection would be relevant for determining which viral mutations might alter the effectiveness of immunity if these individuals were re-exposed to a distinct SARS-CoV-2 variant in the future.

We thus performed escape mapping for samples collected at later time points (76–121 days post-symptom onset) from all 11 individuals for whom we had characterized serum antibody binding at the ~1 month time point. For some but not all individuals there were changes over time in how binding was affected by RBD mutations (**Figures 4, S3, S4** and interactive visualizations at https://jbloomlab.github.io/SARS-CoV-2-RBD_MAP_HAARVI_sera)(Hilton et al., 2020).

**Figure 4.**
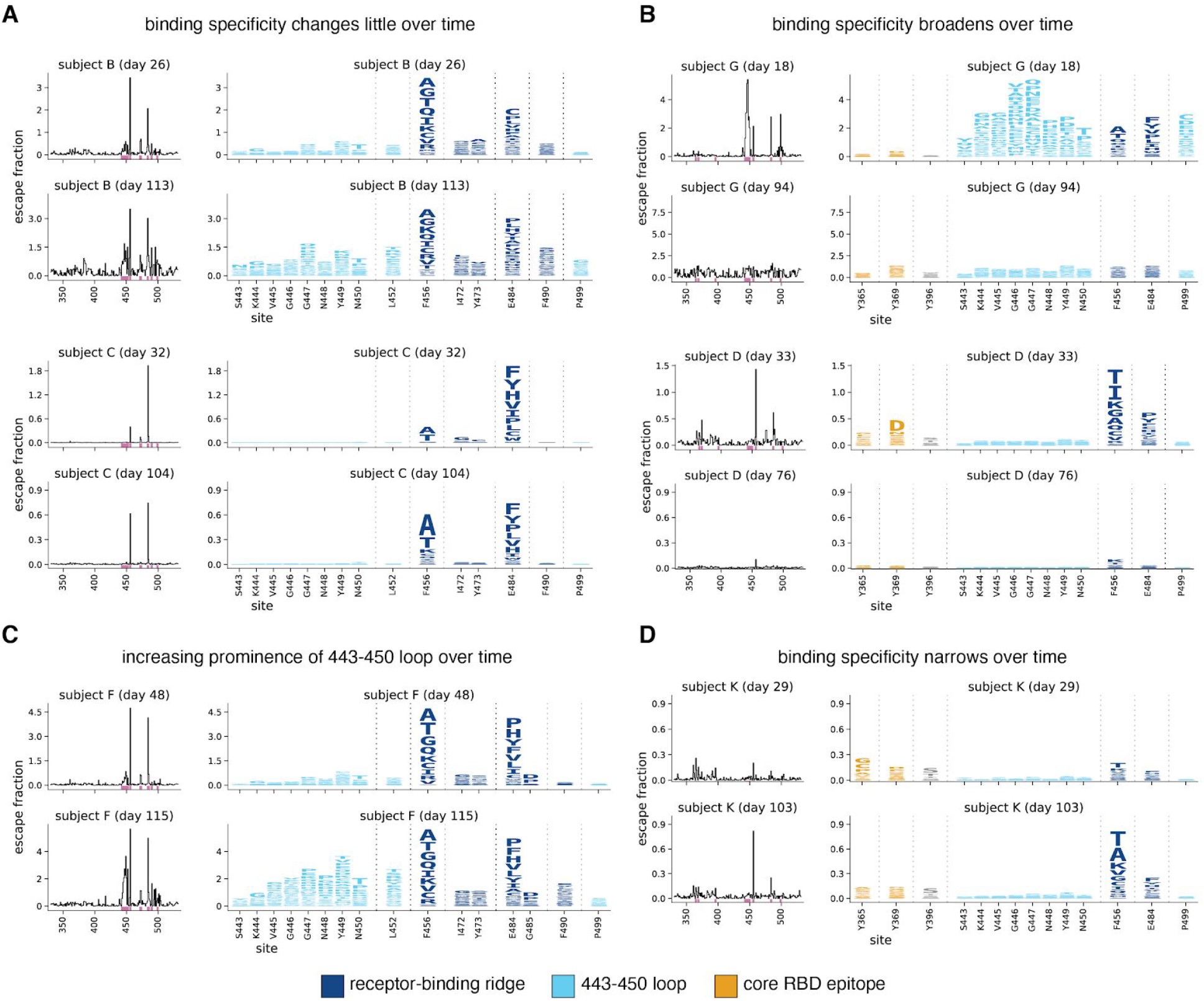
The RBD mutations that affect serum antibody binding change over time for some individuals. Escape maps, colored as in **Figure 2**, demonstrating temporal patterns: **(A)** no change over time, **(B)** broadening over time, **(C)** increasing prominence of one antigenic region, the 443–450 loop, or **(D)** narrowing over time. This figure shows the escape maps over time for 6 of the 11 individuals to illustrate representative trends; see **Figure S3** for escape maps for all individuals at all time points. **Figure S4** shows the effects of mutations at each site projected onto the RBD structure. Different sets of sites are shown in the logo plots in panels **A** and **C**, and in panels **B** and **D**. Sites highlighted in the logo plots are indicated in purple on the x-axes of the associated line plots. The y-axis limits were set as in **Figure 2A** (see **Methods**). Interactive versions of these visualizations are available at https://jbloomlab.github.io/SARS-CoV-2-RBD_MAP_HAARVI_sera/.

Specifically, for over half of the 11 individuals there was little change in which RBD mutations affected serum antibody binding (**Figures 4A, S3, S4**). For two individuals, antibody binding became more broad and less affected by any single RBD mutation (**Figures 4B, S3, S4**). For one individual, mutations in the 443–450 loop had a much larger effect on binding by serum antibodies from the later time compared to the earlier time (**Figure 4C, S3, S4**). Finally, for one individual, there was a strong narrowing of the response, with no single RBD mutation having a large effect on binding by sera from the early time point, but mutations at F456 and to a lesser extent E484 having large effects at the later time point (**Figure 4D, S3, S4**). In summary, while the specificity of serum antibody binding is often maintained over time, in some individuals the specificity broadens to become relatively unaffected by any single RBD mutation, while in other individuals the specificity narrows so that single mutations have a greater impact.

However, there was no clear relationship between changes in the fine specificity of antibody binding and overall serum neutralizing activity. For instance, subject B and subject C maintained similar binding specificities over time (**Figure 4A**), even though the neutralization titers of both subjects’ sera decreased (**Figure 1B**). Similarly, subject D showed major changes in binding specificity over time (**Figure 4B**), although this change in specificity was not accompanied by a substantial change in overall serum neutralization titer.

### For some sera, RBD mutations that reduce antibody binding strongly reduce neutralization

To determine how mutations that reduced serum antibody binding to the RBD affected viral neutralization, we characterized a subset of mutations in neutralization assays with spike-pseudotyped lentiviral particles. For these assays, we chose mutations that our mapping showed had substantial effects on serum antibody binding by samples from multiple individuals, and prioritized mutations present in circulating isolates of SARS-CoV-2.

In many cases, single mutations that were mapped to strongly reduce serum antibody binding also greatly reduced viral neutralization. The effect of mutations at site E484 were particularly striking (**Figure 5A**,**B**). For several sera, the neutralization titer dropped by over an order of magnitude against viruses carrying spikes with E484 mutated to K, Q, or P. For instance, these three mutations to E484 caused 35- to 60-fold decreases in the neutralization titer of the sample collected from subject C at day 32 (**Figure 5A**,**D**). As another example, both E484K and E484Q reduced neutralization by the serum from subject B (day 26) by 10-fold, the same reduction achieved by depleting the serum of all RBD-binding antibodies (**Figure 5A**,**D**).

**Figure 5.**
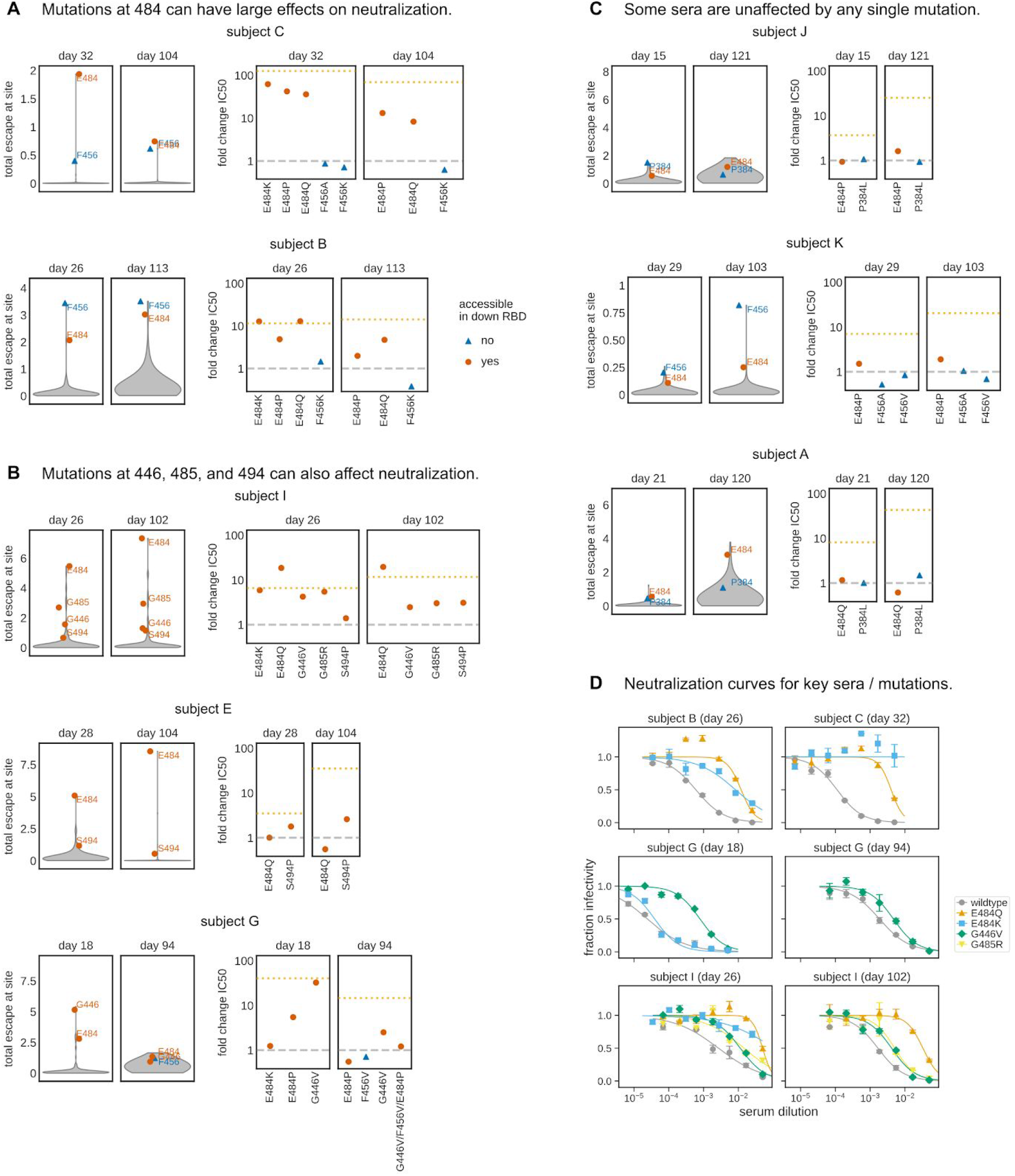
Mutations mapped to reduce serum antibody binding often reduce viral neutralization. **(A)**-**(C)** Violin plots at left show the distribution of how mutations at all sites in the RBD affect serum binding in the mapping experiments. The plots at right then show the effects of tested mutations on neutralization (the fold-change in neutralization inhibitory concentration 50% (IC50)). For instance, the top row in **(A)** shows that mutations at E484 and F456 are mapped to reduce serum antibody binding for subject C at both day 32 and day 104, and that multiple different mutations at E484 but not F456 greatly reduced serum neutralization (e.g., a nearly 100-fold increase in IC50 for E484K for the day 32 serum). Sites that are accessible in the down conformation of the RBD in the context of full spike are indicated by red circles (e.g., E484), and sites that are inaccessible in the RBD’s down conformation are indicated by blue triangles, (e.g., F456). In the plots showing the fold change in IC50s, the dashed gray line indicates a value of one (no change in neutralization), and the dotted orange line indicates the change in inhibitory concentration if all RBD-binding antibodies are removed (see **Figure 1B**). **(D)** Full neutralization curves for a subset of sera and viral mutants demonstrating how E484Q, E484K, G446V, and G485R substantially reduce viral neutralization for some sera. For all neutralization curves used to determine changes in neutralization plotted in **(A)**-**(C)**, see **Figure S5**. The y-axis limits in the violin plots are set as the maximum of the y-axis limit for all time points of a subject in the escape maps in **Figure 2A** and **S3**.

While mutations at E484 generally caused the largest drops in neutralization, other mutations mapped to decrease antibody binding for specific sera also affected neutralization. A dramatic example was G446V, which caused a ~30-fold decrease in the neutralization titer of subject G at day 18 (**Figure 5B**,**D**). Mutations G485R and S494P also caused lesser but still appreciable (~3 to 5-fold) decreases in neutralization titer for a few sera (**Figure 5B**).

In general, there was good concordance between the mapping of how mutations affected serum antibody binding and their impact on viral neutralization. This concordance can be seen in **Figure 5A-C**, where the violin plots show the distributions of the effects of mutations on serum antibody binding across all sites. The sites in the upper tails of these violin plots are ones where mutations had large effects on binding, and mutations to such sites usually reduced neutralization. The one major exception was site F456, where mutations often caused large reductions in binding but never appreciably affected neutralization (**Figure 2A, 5A-C**). This discrepancy between the binding and neutralization effects of mutations is not because antibodies targeting this region are inherently non-neutralizing or unaffected by mutations at site 456, as F456A and F456K disrupt neutralization by two monoclonal antibodies with epitopes that include F456 (Greaney et al., 2020; Shi et al., 2020; Starr et al., 2020a; Zost et al., 2020a) (**Figure S5B**). Rather, we hypothesize that the discrepancy is because we mapped how mutations affected binding using isolated RBD, but in virus the RBD is in the context of full spike, where it can be positioned in either a “down” or “up” conformation (Walls et al., 2020; Wrapp et al., 2020). All sites where mutations that reduced binding also affected neutralization are accessible in both the “down” and “up” conformations, but F456 is only accessible in the “up” conformation. Because the RBD is usually in the “down” conformation (Cai et al., 2020; Ke et al., 2020; Walls et al., 2020), we speculate that while sites accessible in this conformation can be targeted by neutralizing antibodies, they may be subdominant in the context of polyclonal serum binding to full spike even if they are dominant when assaying binding to isolated RBD.

The neutralization assays also validated one of the most notable findings from the mapping: that the antigenic effects of mutations varied markedly across samples from different individuals. For several samples, the maps of binding escape were relatively “flat” with no mutations having disproportionately large effects, and for these samples no tested mutations substantially affected neutralization (**Figure 5C**). Additionally, sometimes the effects of mutations on neutralization changed over time for the same individual. Such a temporal change was especially notable for subject G: mapping of the day 18 sample showed a strong effect of mutations centered around G446, but by day 94 the escape map had flattened (**Figure 4B**). Concordant with the maps, G446V greatly decreased neutralization by subject G’s day 18 sample, but had only a modest effect on the day 94 sample, even when combined with mutations at several other key sites (**Figure 5B**,**D**). These facts highlight how the antigenic effects of mutations vary across people and time, and suggest that some sera are more resistant than others to erosion by viral evolution.

### RBD mutations that reduce serum binding and neutralization in circulating SARS-CoV-2 isolates

To determine the extent that mutations we mapped to affect serum binding are present among circulating SARS-CoV-2 isolates, we determined the frequency of mutations at each RBD site among all SARS-CoV-sequences in GISAID as of Dec-23-2020. We then compared these frequencies to the effects of mutations at each site on serum antibody binding, averaged across all samples (**Figure 6A**).

**Figure 6.**
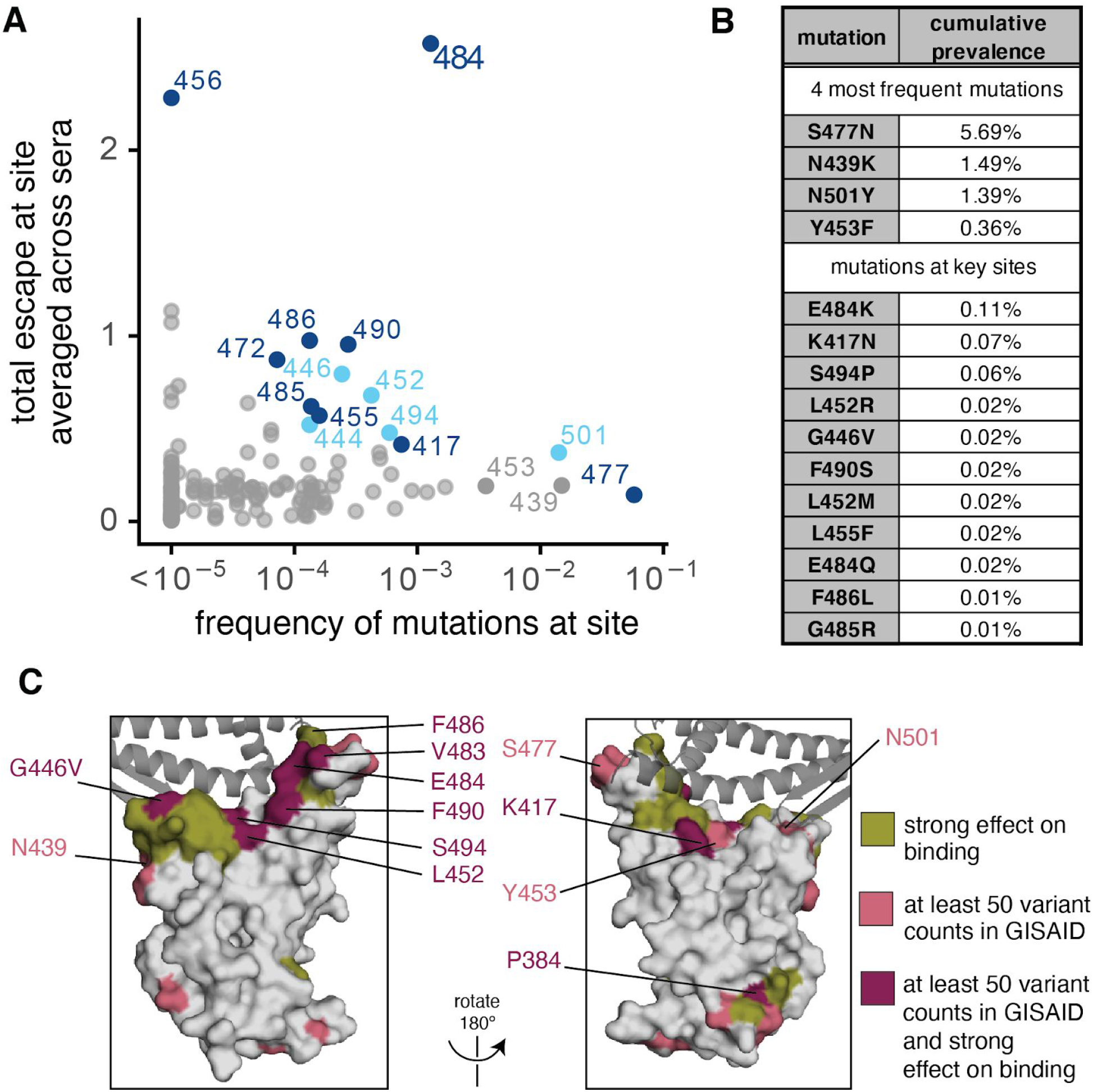
Frequencies of mutations that affect serum antibody binding among circulating SARS-CoV-2 isolates. **(A)** Effects of mutations at each RBD site on serum antibody binding versus frequency of mutations at each site among all SARS-CoV-2 sequences in GISAID as of Dec. 23, 2020. Key sites (see **Methods**) are labeled and colored according to epitope region as in **Figure 2. (B)** Cumulative prevalence for the 4 most frequent mutations and also any mutations at sites labeled in **(A)** with at least 10 counts in GISAID. **(C)** Surface representations of the RBD (PDB 6M0J). Sites where mutations have a strong effect on binding, have circulating variation with >50 total counts in GISAID, or both, are colored in olive, pink, or maroon, respectively. See **Methods** for precise description of highlighted sites. ACE2 is shown as a dark gray cartoon.

It is immediately apparent that the most concerning site of mutations is E484 (**Figure 6A**). E484 is the site where mutations tend to have the largest effect on serum antibody binding to the RBD, and our neutralization assays (**Figure 5A**,**D**) and similar experiments by others (Andreano et al., 2020; Liu et al., 2020b; Weisblum et al., 2020) show that mutations to site E484 reduce the neutralization potency of some human sera by >10 fold, although other sera are unaffected by mutations at this site. Over 0.1% of all sequenced isolates have mutations at this site. Of special note, E484K is present in recently described lineages in South Africa (S.501Y.V2) and Brazil (descended from the B.1.1.28 lineage) (Tegally et al., 2020; Voloch et al., 2020); another mutation at the same site (E484Q) has also been found in a smaller number of human isolates (**Figure 6B**). Two other RBD mutations, K417N and N501Y, co-occur with E484K in the S.501Y.V2 South African lineage (Tegally et al., 2020). K417N escapes neutralization by some monoclonal antibodies (Greaney et al., 2020; Starr et al., 2020a), but mutations to site 417 only modestly affected binding by a few of the samples we assayed (the largest effects were for the last time point for subjects A and J, see **Figure S3**). N501Y increases affinity for ACE2 (Starr et al., 2020b) and is also present in a recently described U.K. lineage (B.1.1.7) that may have increased transmissibility (Kemp et al., 2020a; Public Health England, 2020; Rambaut et al., 2020). Although other mutations at N501 have modest effects on binding by some monoclonal antibodies (Greaney et al., 2020; Starr et al., 2020a), mutations at N501 do not strongly affect binding by any sera we tested (**Figures 2A, S3**).

Several other sites where we mapped mutations to affect serum antibody binding for a few samples also have low-level variation (<0.1%) among circulating viruses (**Figure 6A-C**). These include site G446, where the G446V mutation reduced neutralization by one sample by >10 fold (**Figure 5B**,**D**). Other key sites with circulating variation where mutations impact binding by some samples are indicated in **Figure 6**. Notably, site F456, where mutations consistently affect serum antibody binding but not neutralization, has little variation among circulating viruses (**Figure 6A**).

The four mutations that have the highest frequency among sequenced viruses (S477N, N439K, N501Y, and Y453F; see **Figure 6B**) do not strongly affect serum antibody binding by any samples we tested. As mentioned above, N501Y increases affinity for ACE2, is present in the B.1.1.7 U.K. lineage (Kemp et al., 2020a), and is in the epitope defined by the “443–450 loop” (**Figure 2B**,**3B**)—but does not impact binding by any samples we tested, a result corroborated by live-virus neutralization assays (Menachery, 2020). Y453F and N439K both also increase affinity for ACE2 (Starr et al., 2020b; Thomson et al., 2020), and both escape some monoclonal antibodies (Baum et al., 2020; Starr et al., 2020a; Thomson et al., 2020) but neither greatly impact serum antibody binding by the samples we tested. Finally, S477N also reduces neutralization by some monoclonal antibodies (Liu et al., 2020b), but did not greatly affect binding by the samples we tested.

In summary, our results suggest that E484 is the site of most concern for viral mutations that impact binding and neutralization by polyclonal serum antibodies targeting the RBD. However, mutations at the other serum antibody epitopes (e.g., the 443–450 loop and residues around 484 such as 455, 485, 486, and 490) are also worth monitoring, since they also have antigenic impacts.

## Discussion

We comprehensively mapped how mutations to the SARS-CoV-2 RBD affected binding by the antibodies in convalescent human serum. One major result is that serum antibody binding is predominantly affected by mutations at just a few dominant epitopes in the RBD. In particular, E484 is the site in the RBD where mutations usually have the largest effect on binding and neutralization—possibly because E484 is often targeted by antibodies that utilize heavy-chain germline genes that are common among anti-SARS-CoV-2 RBD antibodies, IGHV3-53 and IGHV3-66 (Barnes et al., 2020a; Greaney et al., 2020; Robbiani et al., 2020; Weisblum et al., 2020; Yuan et al., 2020; Zost et al., 2020b). Mutations at other structurally adjacent sites in the RBD’s receptor binding ridge (e.g., L455, F456, G485, F486, and F490) can also have substantial antigenic effects. Another major epitope centered on the loop formed by residues 443–450 in the RBD’s receptor-binding motif, and mutations in this epitope sometimes strongly affect serum antibody neutralization. A third epitope is in the core of the RBD distal from the receptor-binding motif, although mutations here tend to have smaller effects on serum antibody binding. Notably, RBD mutations reported by other studies to have large effects on serum neutralization are also in the epitope centered around E484 or in the 443–450 loop (Andreano et al., 2020; Li et al., 2020; Weisblum et al., 2020).

While the major serum epitopes are targeted by many well-characterized monoclonal antibodies (Barnes et al., 2020a; Baum et al., 2020; Greaney et al., 2020; Hansen et al., 2020; Starr et al., 2020a), there are also sites where mutations that escape monoclonal antibodies have little effect on serum antibody binding for any sample we tested. For instance, mutations in the S309 epitope footprint (Pinto et al., 2020) and at sites of escape from antibody C135 (e.g., R346 and N440) (Barnes et al., 2020a; Weisblum et al., 2020) had minimal effects on serum antibody binding (**Figures 3, S3**). This lack of concordance between the epitopes of serum and monoclonal antibodies is consistent with other studies reporting that the specificities of potent monoclonal antibodies often do not recapitulate the sera from which they were isolated (Barnes et al., 2020b; Weisblum et al., 2020). These antibodies may be rare in polyclonal sera or the epitopes they target may be subdominant (Piccoli et al., 2020). However, these subdominant epitopes may become more important as SARS-CoV-2 evolves: after mutations at immunodominant sites such as E484 partially erode serum antibody neutralization, the remaining neutralization is presumably due to antibodies targeting previously subdominant epitopes.

Another key finding is that there is extensive person-to-person variation in how mutations affect serum antibody binding and neutralization. For instance, the neutralizing activity of several samples was reduced by >10-fold by single mutations to site E484, but a few samples were essentially unaffected by E484 mutations. Similarly, mutations at sites in the 443–450 loop (e.g., G446V) caused a large drop in serum antibody binding and neutralization for some samples, but had little effect on others. This inter-individual heterogeneity is further compounded by the fact that the effects of mutations sometimes changed over time for samples longitudinally collected from the same individual. These temporal changes could be due to a disproportionate decay in one dominant antibody clonotype, or a relative increase in antibodies targeting other epitopes (Gaebler et al., 2020).

There are several limitations to our study. Most importantly, we only examined mutations to the RBD. While we and others (Piccoli et al., 2020; Steffen et al., 2020) have shown that RBD-binding antibodies contribute the majority of the serum neutralizing activity of most convalescent human sera and plasma, antibodies also target other regions of the spike. For example, mutations and deletions in the NTD can affect serum antibody neutralization (Andreano et al., 2020; Kemp et al., 2020b; Liu et al., 2020a; McCarthy et al., 2020; Voss et al., 2020), and are certainly of great importance. In addition, we only mapped samples from 11 individuals at two time points. Given the substantial inter- and intra-individual heterogeneity, mapping more samples may identify additional sites of importance. On a technical level, we assayed binding of antibodies to isolated RBD expressed by yeast, which implies several limitations. First, we are unable to map the effects of mutations that alter the spike’s overall conformation or affect antibodies spanning quaternary epitopes (Barnes et al., 2020a). Second, our mapping likely overestimates the contributions of antibodies that bind epitopes that are more accessible on isolated RBD than in the context of full spike (e.g., F456). Finally, the N-linked glycans on yeast-expressed proteins are more mannose-rich than those on mammalian-expressed proteins (Hamilton et al., 2003), which could affect measurements of how N-linked glycans affect antibody binding. However, the general consistency of our mapping with our pseudovirus neutralization assays and the serum-escape mutations reported by others suggest that our study successfully defines the major RBD epitopes of convalescent human serum antibodies.

The comprehensive nature of our mapping makes it possible to begin to assess which circulating RBD mutations are likely to have the greatest impact on human immunity. In particular, emerging lineages in South Africa and Brazil carrying the E484K mutation will have greatly reduced susceptibility to neutralization by the polyclonal serum antibodies of some individuals. In contrast, the N501Y mutation present in the U.K. lineage is unlikely to greatly affect neutralization by most human sera, although it could contribute to increased viral titer or enhanced transmissibility (Kidd et al., 2020; Public Health England, 2020). The NTD deletions in this lineage, however, may have an antigenic effect (Andreano et al., 2020; Kemp et al., 2020b; McCarthy et al., 2020). More generally, our serum antibody mapping can be used with other functional characterization to assess the likely antigenic impacts of additional viral mutations that emerge in the future.

Our mapping also reveals broader features of antibody immunity that are relevant to SARS-CoV-2 evolution. One reason that influenza virus undergoes such rapid antigenic evolution is that neutralizing human immunity often focuses on just a few residues in hemagglutinin, such that a single mutation can dramatically reduce neutralization (Lee et al., 2019). In contrast, while measles virus can escape neutralization by monoclonal antibodies, polyclonal serum targets multiple co-dominant measles epitopes, meaning that no single mutation has a large effect on neutralization (Muñoz-Alí a et al., 2020). Our results show that polyclonal antibody immunity to the SARS-CoV-2 RBD is sometimes focused as for influenza, but in other cases more broadly targets the RBD in a way that mitigates the effect of any single mutation. This heterogeneity in the antigenic impacts of RBD mutations implies that the immunity of different individuals will be impacted differently by viral evolution. It also suggests that an important area for future work is understanding how viral mutations impact vaccine-elicited immunity, and using this knowledge to design vaccines that are robust to viral antigenic evolution.

## Acknowledgments

We thank Adam Dingens for experimental assistance; the Flow Cytometry and Genomics core facilities at the Fred Hutchinson Cancer Research Center for experimental support, especially Dolores Covarrubias and Andy Marty; Neil King, Alexandra Walls, David Veesler, and the UW Institute for Protein Design for purified RBD and spike proteins; and Seth Zost and James Crowe of Vanderbilt University for monoclonal antibodies. We also thank all research participants in the Hospitalized or Ambulatory Adults with Respiratory Viral Infections (HAARVI) study for their generous participation and all HAARVI study researchers and staff, especially Caitlin Wolf. This work was supported by the NIAID / NIH (R01AI141707 and R01AI127893 to J.D.B., T32AI083203 to A.J.G., and F30AI149928 to K.H.D.C.) and the Gates Foundation (INV-004949). T.N.S. is a Washington Research Foundation Innovation Fellow at the University of Washington Institute for Protein Design and a Howard Hughes Medical Institute Fellow of the Damon Runyon Cancer Research Foundation (DRG-2381-19). J.D.B. is an Investigator of the Howard Hughes Medical Institute. The content is solely the responsibility of the authors and does not necessarily represent the official views of the US government or the other sponsors.

## Author contributions

Conceptualization, A.J.G., H.Y.C., and J.D.B.; Methodology, A.J.G., K.H.D.C., T.N.S., and J.D.B.; Investigation, A.J.G., A.L., K.H.D.C., and K.M.; Code, A.J.G., T.N.S., and J.D.B.; Formal Analysis, A.J.G. and J.D.B.; Validation, A.J.G., A.N.L., and K.H.D.C.; Resources, H.Y.C.; Writing – Original Draft, A.J.G. and J.D.B.; Writing – Review and Editing, all authors; Supervision, H.Y.C. and J.D.B.

## Declarations of Interests

H.Y.C. is a consultant for Merck and Pfizer and receives research funds from Cepheid, Ellume, Genentech, and Sanofi-Pasteur.

## Methods

### Data and code availability

We provide data and code in the following ways:

- The complete code for the full computational data analysis pipeline of the mapping experiments is available on GitHub at https://github.com/jbloomlab/SARS-CoV-2-RBD_MAP_HAARVI_sera
- A Markdown summary of the analysis workflow with renderings of all the code is on GitHub at https://github.com/jbloomlab/SARS-CoV-2-RBD_MAP_HAARVI_sera/blob/main/results/summary/summary.md
- The escape fraction measured for each mutation in **Supplementary Table 3** and also at https://github.com/jbloomlab/SARS-CoV-2-RBD_MAP_HAARVI_sera/blob/main/results/escape_scores/escape_fracs.csv
- The barcode counts for each RBD variant in each mapping condition are at https://github.com/jbloomlab/SARS-CoV-2-RBD_MAP_HAARVI_sera/blob/main/results/counts/variant_counts.csv
- All raw sequencing data are available on the NCBI Short Read Archive at BioProject PRJNA639956 (https://www.ncbi.nlm.nih.gov/bioproject/PRJNA639956), BioSample SAMN17185313 (https://www.ncbi.nlm.nih.gov/biosample/?term=SAMN17185313).

### SARS-CoV-2 convalescent human sera

Plasma samples were previously described (Crawford et al., 2020a) and collected as part of a prospective longitudinal cohort study of individuals with SARS-CoV-2 infection in Seattle, WA February-July 2020. See **Supplementary Table 1** for the sample metadata, which is also described in (Crawford et al., 2020a). That table also links the sample IDs used in (Crawford et al., 2020a) to the names used for the sera in this paper. All sera were heat-inactivated prior to use by treatment at 56 C for 60 minutes. Prior to use in each assay, plasma samples were centrifuged for 15 min at 2000 x*g* to pellet platelets.

### RBD deep mutational scanning library

The yeast-display RBD mutant libraries are previously described (Greaney et al., 2020; Starr et al., 2020b). Briefly, duplicate mutant libraries were constructed in the spike receptor binding domain (RBD) from SARS-CoV-2 (isolate Wuhan-Hu-1, Genbank accession number MN908947, residues N331-T531) and contain 3,804 of the 3,819 possible amino-acid mutations, with >95% present as single mutants. Each RBD variant was linked to a unique 16-nucleotide barcode sequence to facilitate downstream sequencing. As previously described, libraries were sorted for RBD expression and ACE2 binding to eliminate RBD variants that are completely misfolded or non-functional (i.e., lacking modest ACE2 binding affinity) (Greaney et al., 2020).

### FACS sorting of yeast libraries to select mutants with reduced binding by polyclonal sera

Serum mapping experiments were performed in biological duplicate using the independent mutant RBD libraries, as previously described for monoclonal antibodies (Greaney et al., 2020), with the following modifications: Mutant yeast libraries induced to express RBD were washed and incubated with serum at a range of dilutions for 1 h at room temperature with gentle agitation. For each serum, we chose a sub-saturating dilution such that the amount of fluorescent signal due to serum antibody binding to RBD was approximately equal across sera. The exact dilution used for each serum is given in **Supplementary Table 2**. After the serum incubations, the libraries were secondarily labeled with 1:100 FITC-conjugated anti-MYC antibody (Immunology Consultants Lab, CYMC-45F) to label for RBD expression and 1:200 Alexa-647- or DyLight-405-conjugated goat anti-human-IgA+IgG+IgM (Jackson ImmunoResearch 109-605-064 or 109-475-064, respectively) to label for bound serum antibodies. A flow cytometric selection gate was drawn to capture 3–6% of the RBD mutants with the lowest amount of serum binding for their degree of RBD expression (**Figure S1A-C**). We also measured what fraction of cells expressing unmutated RBD fell into this gate when stained with 1x and 0.1x the concentration of serum. For each sample, approximately 10 million RBD+ cells (range 7.4e6 to 1.7e7 cells) were processed on the cytometer, with between 2e5 and 8e5 serum-escaped cells collected per sample (see percentages in **Supplementary Table 2**). Antibody-escaped cells were grown overnight in SD-CAA (6.7g/L Yeast Nitrogen Base, 5.0g/L Casamino acids, 1.065 g/L MES acid, and 2% w/v dextrose) to expand cells prior to plasmid extraction.

### DNA extraction and Illumina sequencing

Plasmid samples were prepared from 30 OD units (1.6e8 cfus) of pre-selection yeast populations and approximately 5 OD units (~3.2e7 cfus) of overnight cultures of serum-escaped cells (Zymoprep Yeast Plasmid Miniprep II) as previously described (Greaney et al., 2020). The 16-nucleotide barcode sequences identifying each RBD variant were amplified by PCR and prepared for Illumina sequencing as described in (Starr et al., 2020b). Barcodes were sequenced on an Illumina HiSeq 3500 with 50 bp single-end reads. To minimize noise from inadequate sequencing coverage, we ensured that each antibody-escape sample had at least 2.5x as many post-filtering sequencing counts as FACS-selected cells, and reference populations had at least 2.5e7 post-filtering sequencing counts.

### Analysis of deep sequencing data to compute each mutation’s serum escape fraction

Escape fractions were computed as described in (Greaney et al., 2020), with minor modifications as noted below. We used the `dms_variants` package (https://jbloomlab.github.io/dms_variants/, version 0.8.2) to process Illumina sequences into counts of each barcoded RBD variant in each pre-sort and antibody-escape population using the barcode/RBD look-up table from (Starr et al., 2020a).

For each serum selection, we computed the “escape fraction” for each barcoded variant using the deep sequencing counts for each variant in the original and serum-escape populations and the total fraction of the library that escaped antibody binding via the formula provided in (Greaney et al., 2020). These escape fractions represent the estimated fraction of cells expressing that specific variant that fall in the serum escape bin, such that a value of 0 means the variant is always bound by serum and a value of 1 means that it always escapes serum binding. We then applied a computational filter to remove variants with low sequencing counts or highly deleterious mutations that might cause antibody escape simply by leading to poor expression of properly folded RBD on the yeast cell surface (Greaney et al., 2020; Starr et al., 2020b). Specifically, we removed variants that had (or contained mutations with) ACE2 binding scores < −2.35 or expression scores < −1, using the variant- and mutation-level deep mutational scanning scores from (Starr et al., 2020b). Note that these filtering criteria are slightly more stringent than those used in (Greaney et al., 2020) but are identical to those used in (Starr et al., 2020a).

We next deconvolved variant-level escape scores into escape fraction estimates for single mutations using global epistasis models (Otwinowski et al., 2018) implemented in the `dms_variants` package, as detailed at (https://jbloomlab.github.io/dms_variants/dms_variants.globalepistasis.html) and described in (Greaney et al., 2020). The reported scores throughout the paper are the average across the libraries; these scores are also in **Supplementary Table 3**. Correlations in final single-mutant escape scores are shown in **Figure S2D-E**.

For plotting and analyses that required identifying RBD sites of “strong escape” (e.g., choosing which sites to show in logo plots in **Figures 2A**,**B, and S4** or label in **Figure 4B**), we considered a site to mediate strong escape if the total escape (sum of mutation-level escape fractions) for that site exceeded the median across sites by >5 fold, and was at least 5% of the maximum for any site. We also included site K417, which did not meet this threshold but was of interest due to its frequency among circulating viruses.

Full documentation of the computational analysis is at https://github.com/jbloomlab/SARS-CoV-2-RBD_MAP_HAARVI_sera.

### Generation of pseudotyped lentiviral particles

We used spike-pseudotyped lentiviral particles that were generated essentially as described in (Crawford et al., 2020b), using a codon-optimized SARS-CoV-2 spike from Wuhan-Hu-1 that contains a 21-amino-acid deletion at the end of the cytoplasmic tail (Crawford et al., 2020a) and the D614G mutation that is now predominant in human SARS-CoV-2 (Korber et al., 2020). The plasmid encoding this spike, HDM_Spikedelta21_D614G, is available from Addgene (#158762), and the full sequence is at (https://www.addgene.org/158762). Point mutations were introduced into the RBD of this plasmid via site-directed mutagenesis.

To generate these spike-pseudotyped lentiviral particles (Crawford et al., 2020b), 6e5 293T cells per well were seeded in 6-well plates in 2 mL D10 growth media (DMEM with 10% heat-inactivated FBS, 2 mM l-glutamine, 100 U/mL penicillin, and 100 μg/mL streptomycin). 24h later, cells were transfected using BioT transfection reagent (Bioland Scientific, Paramount, CA, USA) with a Luciferase_IRES_ZsGreen backbone, Gag/Pol lentiviral helper plasmid, and wildtype or mutant SARS-CoV-2 spike plasmids. Media was changed to fresh D10 at 24 h post-transfection. At 60 h post-transfection, viral supernatants were collected, filtered through a 0.45 μm SFCA low protein-binding filter, and stored at −80°C.

### Neutralization assays

293T-ACE2 cells (BEI NR-52511) were seeded at 1.25e4 cells per well in 50 μL D10 in poly-L-lysine coated, black-walled, 96-well plates (Greiner 655930). 24 h later, pseudotyped lentivirus supernatants were diluted to ~200,000 relative luciferase units (RLU) per well (determined by titering as previously described (Crawford et al., 2020b)) and incubated with a range of dilutions of sera for 1 h at 37 °C. 100 μL of the virus-antibody mixture was then added to cells.

At ~70 h post-infection, luciferase activity was measured using the Bright-Glo Luciferase Assay System (Promega, E2610). Fraction infectivity of each serum antibody-containing well was calculated relative to a “no-serum” well inoculated with the same initial viral supernatant (containing wildtype or mutant RBD) in the same row of the plate. We used the `neutcurve` package (https://jbloomlab.github.io/neutcurve) to calculate the inhibitory concentration 50% (IC50) and the neutralization titer 50% (NT50), which is simply 1/IC50, of each serum against each virus by fitting a Hill curve with the bottom fixed at 0 and the top fixed at 1. The full neutralization curves are in **Figure S5**.

### Depletion of RBD-binding antibodies from polyclonal sera

Magnetic beads conjugated to the SARS-CoV-2 RBD (AcroBiosystems, MBS-K002) were prepared according to the manufacturer’s protocol. Beads were resuspended in ultrapure water at 1 mg beads/mL and a magnet was used to wash the beads 3 times in PBS with 0.05% BSA. Beads were then resuspended in PBS with 0.05% BSA at 1 mg beads /mL. Beads (manufacturer-reported binding capacity of 10-40 ug/mL anti-RBD antibodies) were incubated with human serum at a 3:1 ratio beads:serum (150 uL beads + 50 uL serum), rotating overnight at 4°C. A magnet was used to separate antibodies that bind RBD from the supernatant, and the supernatant (the post-RBD antibody depletion sample) was removed. A mock depletion (pre-depletion sample) was performed by adding 150 uL of PBS + 0.05% BSA and incubating rotating overnight at 4°C. For the neutralization assays on these sera depleted of RBD-binding antibodies shown in **Figure S1C**; the reported serum dilution is corrected for the dilution incurred by the depletion process.

### Measurement of serum binding to RBD or spike by ELISA

The IgG ELISAs for spike protein and RBD were conducted as previously described (Dingens et al., 2020). Briefly, ELISA plates were coated with recombinant spike and RBD antigens described in (Dingens et al., 2020) at 2 μg/mL. Five 3-fold serial dilutions of sera beginning at 1:100 were performed in phosphate-buffered saline with 0.1% Tween nonfat dry milk. Dilution series of the “synthetic” sera comprised of the anti-RBD antibody rREGN10987 (Hansen et al., 2020) or anti-NTD antibody 4A8 (Chi et al., 2020) and pooled pre-pandemic human sera from 2017-2018 (Gemini Biosciences; nos. 100–110, lot H86W03J; pooled from 75 donors) were performed such that the anti-spike antibody was present at a highest concentration of 0.25 μg/mL. Both antibodies were recombinantly produced by Genscript. The rREGN10987 is that used in (Starr et al., 2020a) and the variable domain heavy and light chain sequences for r4A8 were obtained from Genbank GI 1864383732 and 1864383733 (Chi et al., 2020) and produced on a human IgG1 and IgK background, respectively. Pre-pandemic serum alone, without anti-RBD antibody depletion, was used as a negative control, averaged over 2 replicates. The area under the curve (AUC) was calculated as the area under the titration curve with the serial dilutions on a log-scale.

### Analysis of RBD mutations among circulating SARS-CoV-2 isolates

All 283,908 spike sequences on GISAID as of Dec-23-2020 were downloaded and aligned via mafft (Katoh and Standley, 2013). Sequences from non-human origins and sequences containing gap characters or excessive mutations were removed, leaving 263,217 sequences. The code that performs this alignment and filtering is at https://github.com/jbloomlab/SARS-CoV-2-RBD_MAP_HAARVI_sera/blob/main/results/summary/gisaid_rbd_mutations.md. The counts and frequencies of mutations at each RBD site were then computed using this filtered sequence set. We acknowledge all GISAID contributors for sharing sequencing data (https://github.com/jbloomlab/SARS-CoV-2-RBD_MAP_HAARVI_sera/blob/main/data/gisaid_hcov-19_acknowledgement_table_2020_12_30.pdf).

Sites and mutations highlighted in **Figure 6** were chosen as follows. Sites in the RBM containing the 4 RBD mutations with the highest cumulative frequency (S477N, N439K, N501Y, and Y453F), the two sites with the highest total escape (F456 and E484), and sites that have >=30 variant counts in GISAID and are sites of strong escape for any serum, are labeled in **Figure 6A**. The labeled sites are colored according to epitope region as in **Figure 2**. Figure 6B highlights the 4 most frequent mutations and also any mutations at the other sites labeled in **Figure 6A** with at least 10 or more counts in GISAID. **Figure 6C** highlights sites where mutations have a strong effect on binding of at least 1 serum or have circulating variation with >50 counts in GISAID. Site K417 was also of interest due to the presence of the K417N mutation in a recently identified lineage (Tegally et al., 2020), and thus is also highlighted in each panel in **Figure 6**.

### Data visualization

The static logo plot visualizations of the escape maps in the paper figures were created using the `dmslogo` package (https://jbloomlab.github.io/dmslogo, version 0.3.2) and in all cases the height of each letter indicates the escape fraction for that amino-acid mutation calculated as described above. For each serum, the y-axis is scaled to be the greatest of (a) the maximum site-wise escape metric observed for that serum, (b) 20x the median site-wise escape fraction observed across all sites for that serum, or (c) an absolute value of 1.0 (to appropriately scale sera that are not “noisy” but for which no mutation has a strong effect on serum binding). Site C361 has been removed from the plots, because while mutations at this site reduce serum binding, these mutations ablate a disulfide bond in the core RBD that is important for proper folding of the RBD and likely result in a grossly misfolded RBD and do not represent specific serum escape mutations. Site K417 has been added to **Figure S3** due to its frequency among circulating viruses. The code that generates these logo plot visualizations is available at https://github.com/jbloomlab/SARS-CoV-2-RBD_MAP_HAARVI_sera/blob/main/results/summary/escape_profiles.md.

In many of the visualizations (e.g., **Figures 2, 4, 6A, S3, and S4**), the RBD sites are categorized by epitope region (core-RBD epitope, receptor-binding ridge, or 443–450 loop) and colored accordingly. We define the core-RBD epitope as residues 365–372+382–386, the receptor-binding ridge epitope to be residues 417+455+456+471–490, and the 443–450 loop epitope to be residues 443–452+494–501.

For the static structural visualizations in the paper figures, the RBD surface (PDB: 6M0J, (Lan et al., 2020)) was colored by the site-wise escape metric at each site, with white indicating no escape and red scaled to be the same maximum used to scale the y-axis in the logo plot escape maps, determined as described above.We created interactive structure-based visualizations of the escape maps using `dms-view` (Hilton et al., 2020) that are available at https://jbloomlab.github.io/SARS-CoV-2-RBD_MAP_HAARVI_sera. The logo plots in these escape maps can be colored according to the deep mutational scanning measurements of how mutations affect ACE2 binding or RBD expression as described above.

## Supplementary Figures

**Figure S1.**
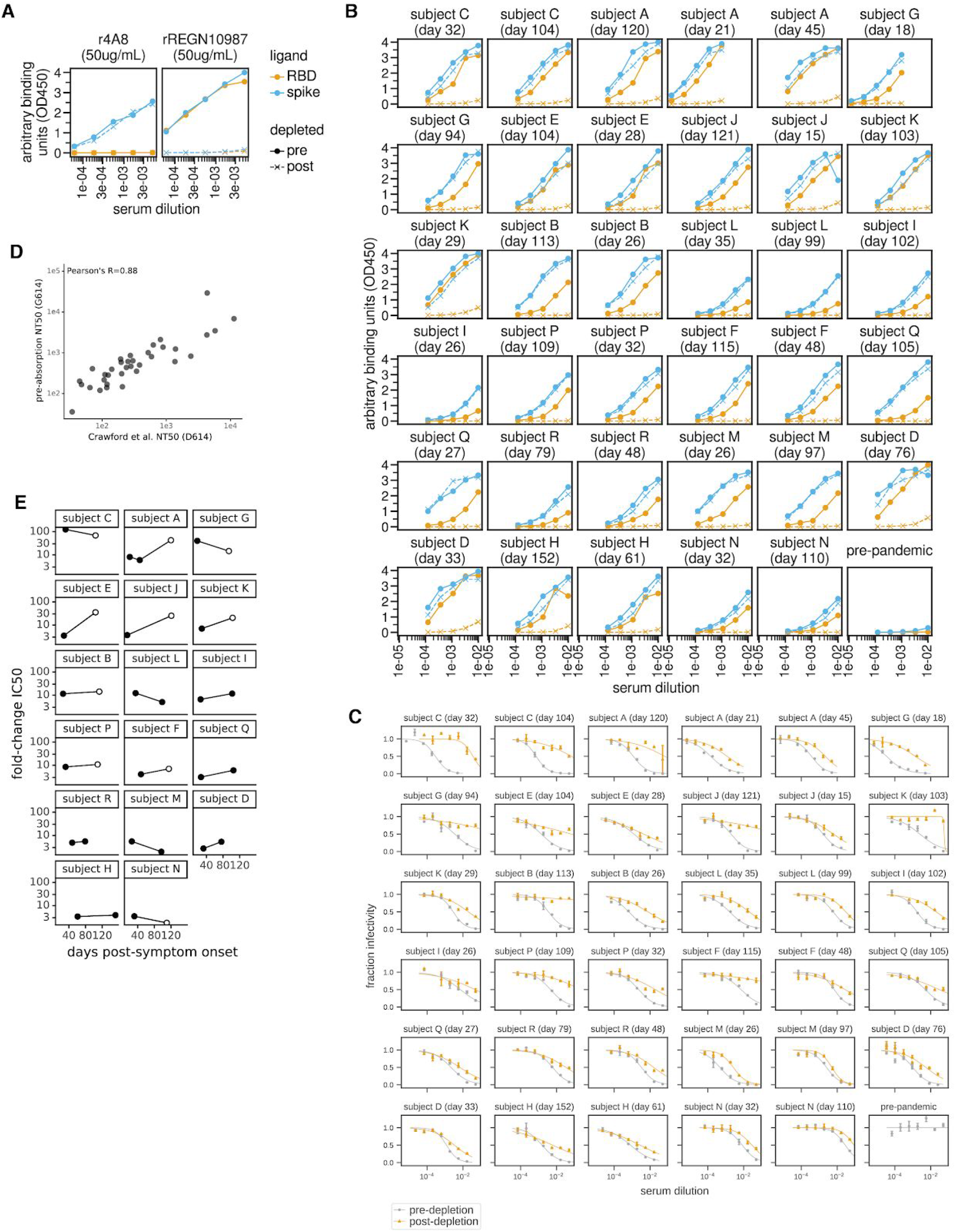
Raw ELISA and neutralization curves of sera pre- and post-depletion of RBD-targeting antibodies, related to Figure 1. **(A)** Effect of RBD antibody depletion on binding to RBD and spike by “synthetic sera” comprised of pre-pandemic pooled serum with the NTD-targeting antibody r4A8 (Chi et al., 2020) or RBD-targeting antibody rREGN10987 (Hansen et al., 2020). Antibodies were added to pre-pandemic serum at 50 μg/mL. The x-axis indicates the dilution factor of the serum+antibody mix, and the y-axis is the ELISA reading at each dilution. **(B)** Raw ELISA binding curves of plasma to RBD and spike before and after depletion of RBD-binding antibodies. Legend for panels **(A)** and **(B)**: orange is RBD binding, blue is spike binding; filled circles with solid lines represent pre-depletion, and x’s with dashed lines represent post-depletion of anti-RBD antibodies. **(C)** Raw neutralization curves for plasma before (gray) and after (orange) depletion of RBD-binding antibodies. Neutralization assays were performed with lentiviruses pseudotyped with spike D614G, the predominant SARS-CoV-2 circulating variant. **(D)** Correlation between previously measured neutralization titers 50% (NT50) with spike D614-spike-pseudotyped lentivirus (Crawford et al., 2020a) and pre-depletion neutralization titers measured with G614-spike-pseudotyped lentivirus (present study), Pearson’s R = 0.88. **(E)** Change in the amount of neutralizing activity that is due to RBD-binding antibodies over time for each individual. Each point gives the fold-change in neutralization inhibitory concentration 50% (IC50) post-versus pre-depletion for plasma isolated at the indicated time, such that larger values indicate that more of the neutralizing activity is due to RBD-binding antibodies. Open circles represent samples for which the post-depletion NT50 was at the limit of detection, i.e., less than 20 (see **Figure 1B**); these circles are therefore lower bounds in the fold-change in IC50.

**Figure S2.**
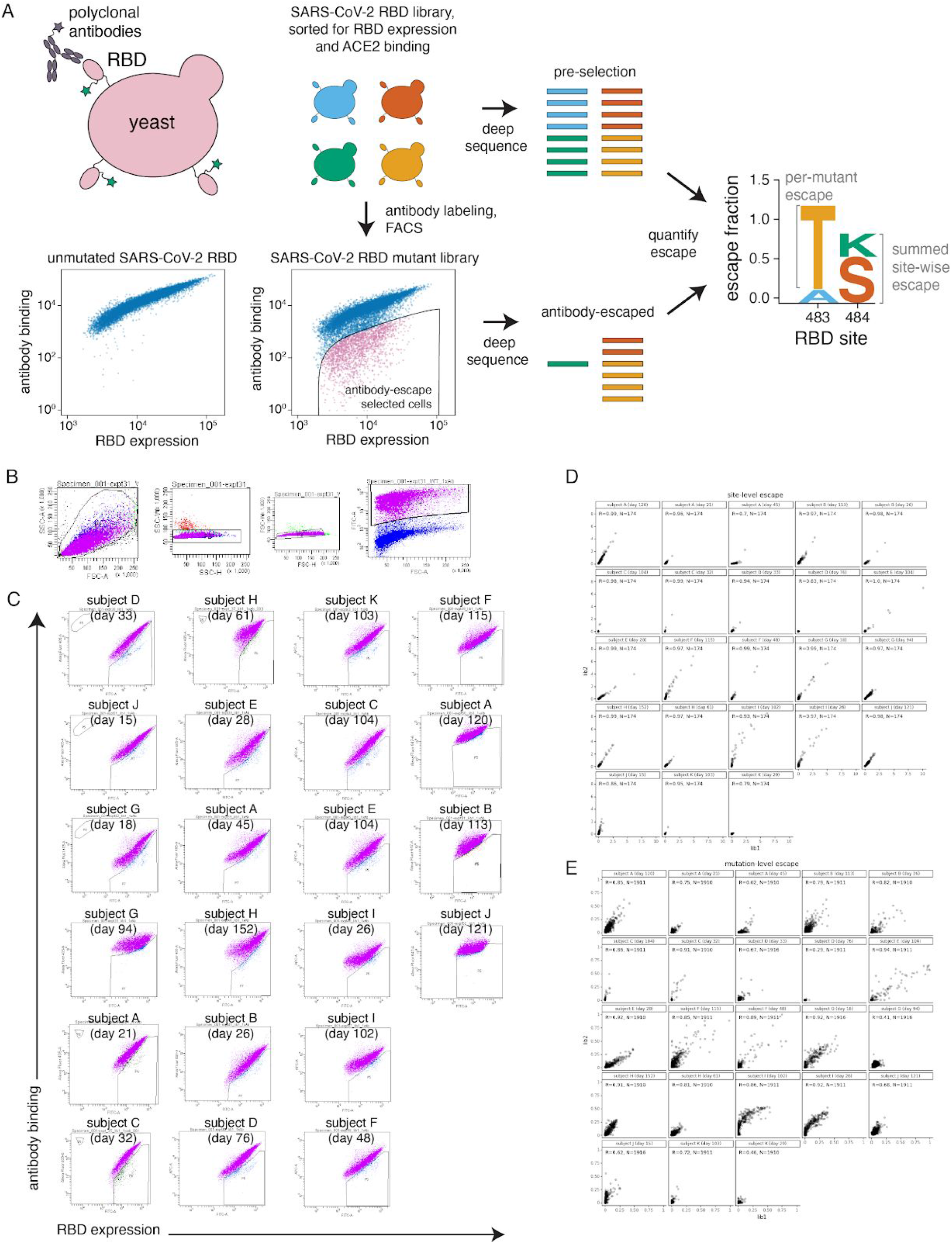
Approach for mapping RBD mutations that reduce binding by polyclonal sera, related to Figure 2. **(A)** The RBD is expressed on the surface of yeast. Flow cytometry can be used to quantify both RBD expression (via a C-terminal MYC tag) and antibody binding to the RBD protein expressed on the surface of each yeast cell. A library of yeast expressing different RBD mutants were incubated with polyclonal serum and serum antibody binding was detected using a IgA+IgG+IgM secondary antibody. We then used FACS to enrich for cells expressing RBD that bound reduced levels of antibody, and used deep sequencing to quantify the frequency of each mutation in the initial and “antibody escape” cell populations. We quantified the effect of each mutation as the “escape fraction,” which represents the fraction of cells expressing RBD with that mutation that fell in the “antibody escape” FACS bin. Escape fractions are represented in logo plots, with the height of each letter proportional to the effect of that amino acid mutation on antibody binding. The site-level escape metric is the sum of the escape fractions of all mutations at a site. Note that both experimental and computational filtering steps were used to remove RBD mutants that were misfolded or completely unable to bind the ACE2 receptor (see **Methods**). **(B)** Representative plots of nested FACS gating strategy used for all serum selection experiments to select for single cells (SSC-A vs. FSC-A, and FSC-W vs. FSC-H) that also express RBD (FITC-A vs. FSC-A). **(C)** FACS gating strategy for one of two independent libraries to select cells expressing RBD mutants with reduced binding by polyclonal sera. Gates were set manually during sorting, aiming for 3-6% of the RBD+ library to fall into the selection gate (cells in blue). The same gate was set for both independent libraries stained with each serum, and the FACS scatter plots looked qualitatively similar between the two libraries. For information on the fraction of library cells that fall into each selection gate, see **Supplementary Table 2. (D)** Correlation plots of site-level escape for each of the two independent RBD mutant libraries for each serum. Site-level escape is the sum of escape fraction for each mutation at a site. **(E)** Correlation plots of mutation-level escape for each of the two independent RBD mutant libraries for each serum.

**Figure S3.**
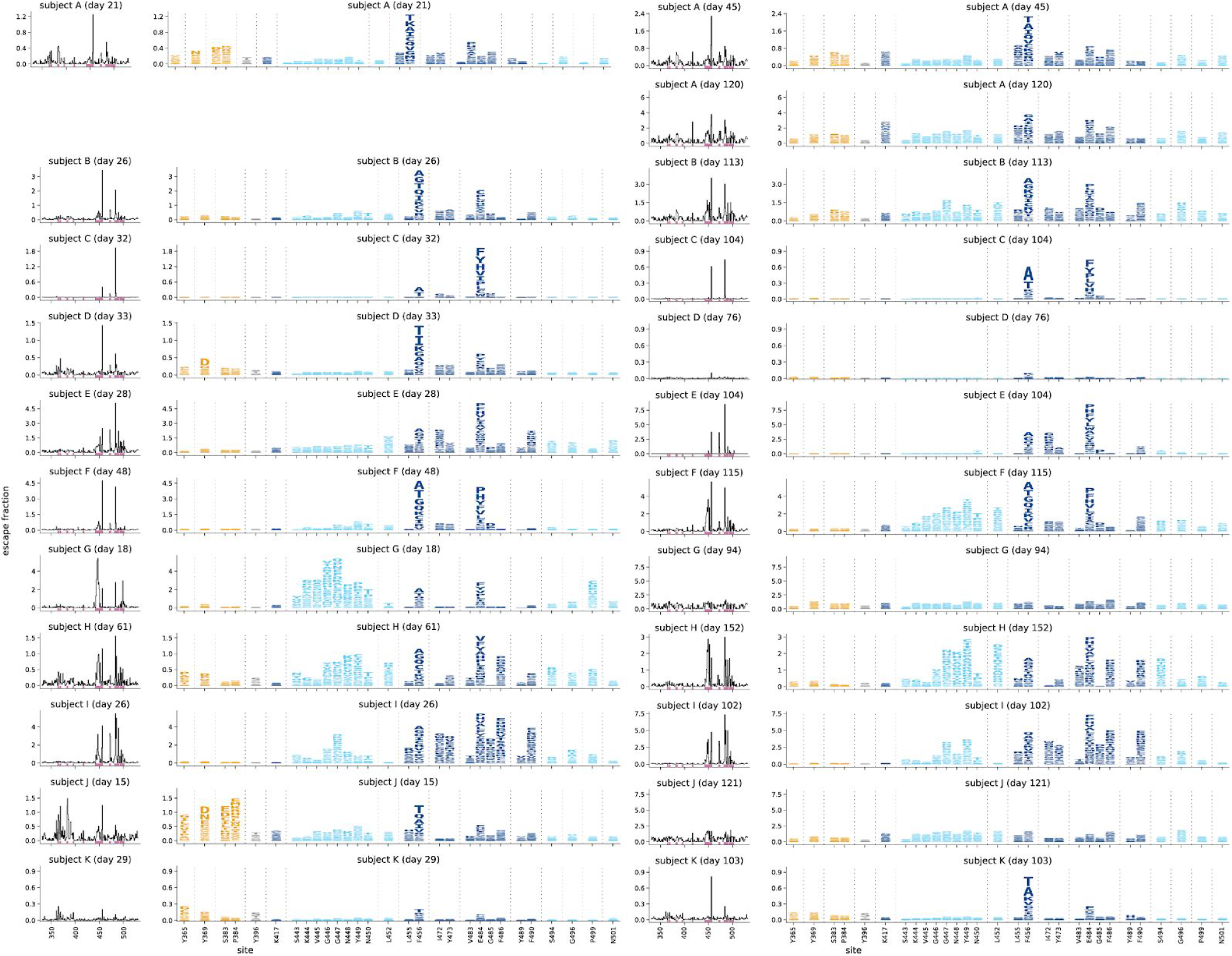
Escape maps over time for all study subjects and time points, related to Figure 4. Escape maps for all individuals and time points, with 2 time points shown side-by-side, ordered as in **Figure 2A**. Escape fractions are comparable across sites within a sample, but not necessarily between samples due to the use of sample-specific FACS gates—therefore, for each sample, the y-axis is scaled independently (see **Methods**). Sites are colored by RBD epitope region as in **Figure 2**. Sites shown in logo plots, highlighted in purple in line plots at left, are sites of strong escape for any of the 23 sera, plus sites K417 and N501. Interactive versions of these escape maps are available at https://jbloomlab.github.io/SARS-CoV-2-RBD_MAP_HAARVI_sera/.

**Figure S4.**
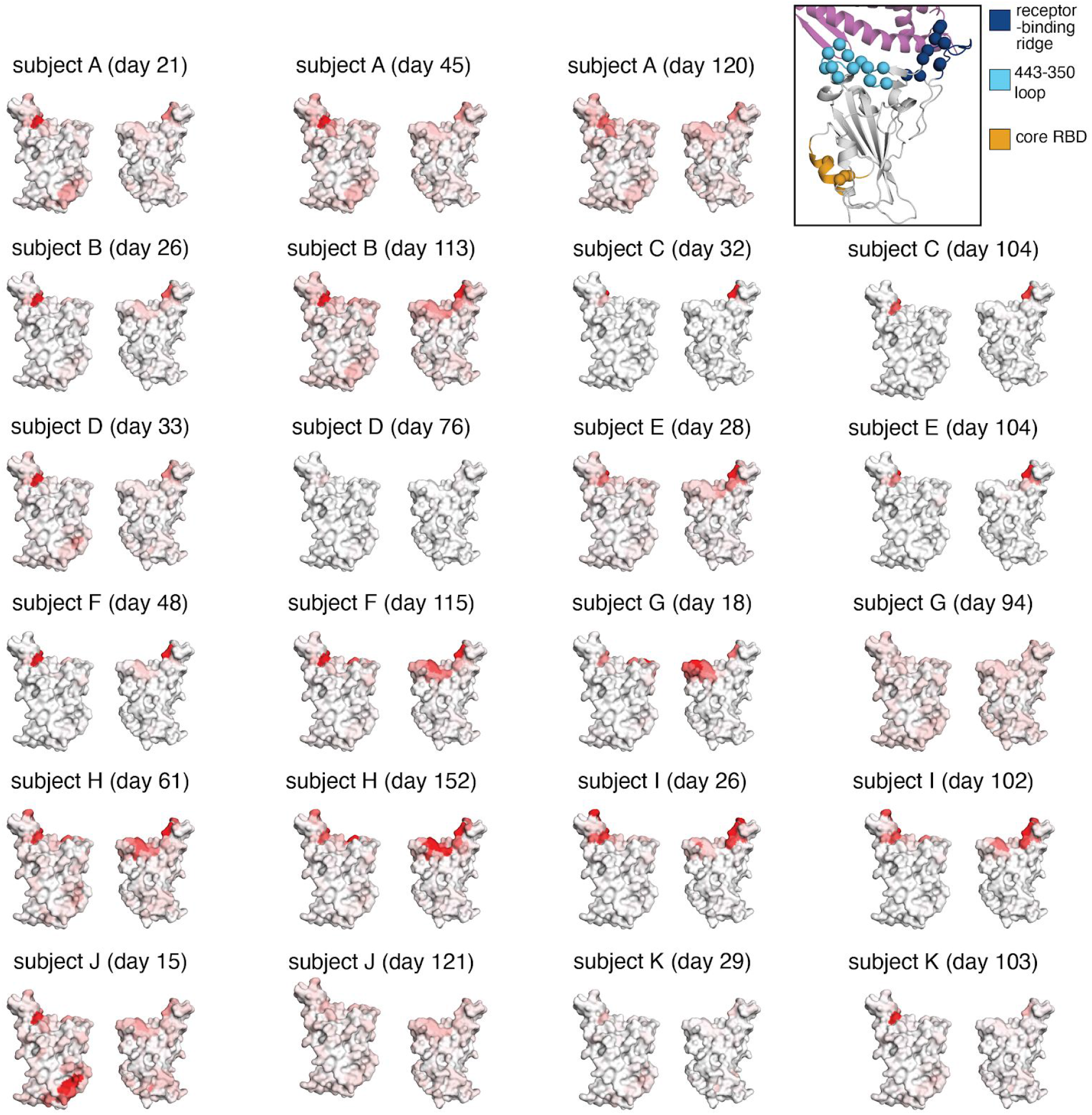
Regions in the RBD where mutations reduce binding by sera antibodies for all study subjects and samples over time, related to Figure 4. The structures show the effects of mutations at each site projected onto the RBD structure using a white-to-red color scale as in **Figure 3A-D**. The color scale for each serum is scaled to span the same range as the y-axis for that serum in **Figure S3**. Top right inset: the alpha-carbon of any site of strong escape (all sites shown in the logo plots in **Figure S3**) is shown as a sphere on a cartoon representation of the RBD (PDB 6M0J). The RBD is colored as in **Figure 2B**. Interactive versions of these structural visualizations are available at https://jbloomlab.github.io/SARS-CoV-2-RBD_MAP_HAARVI_sera/.

**Figure S5.**
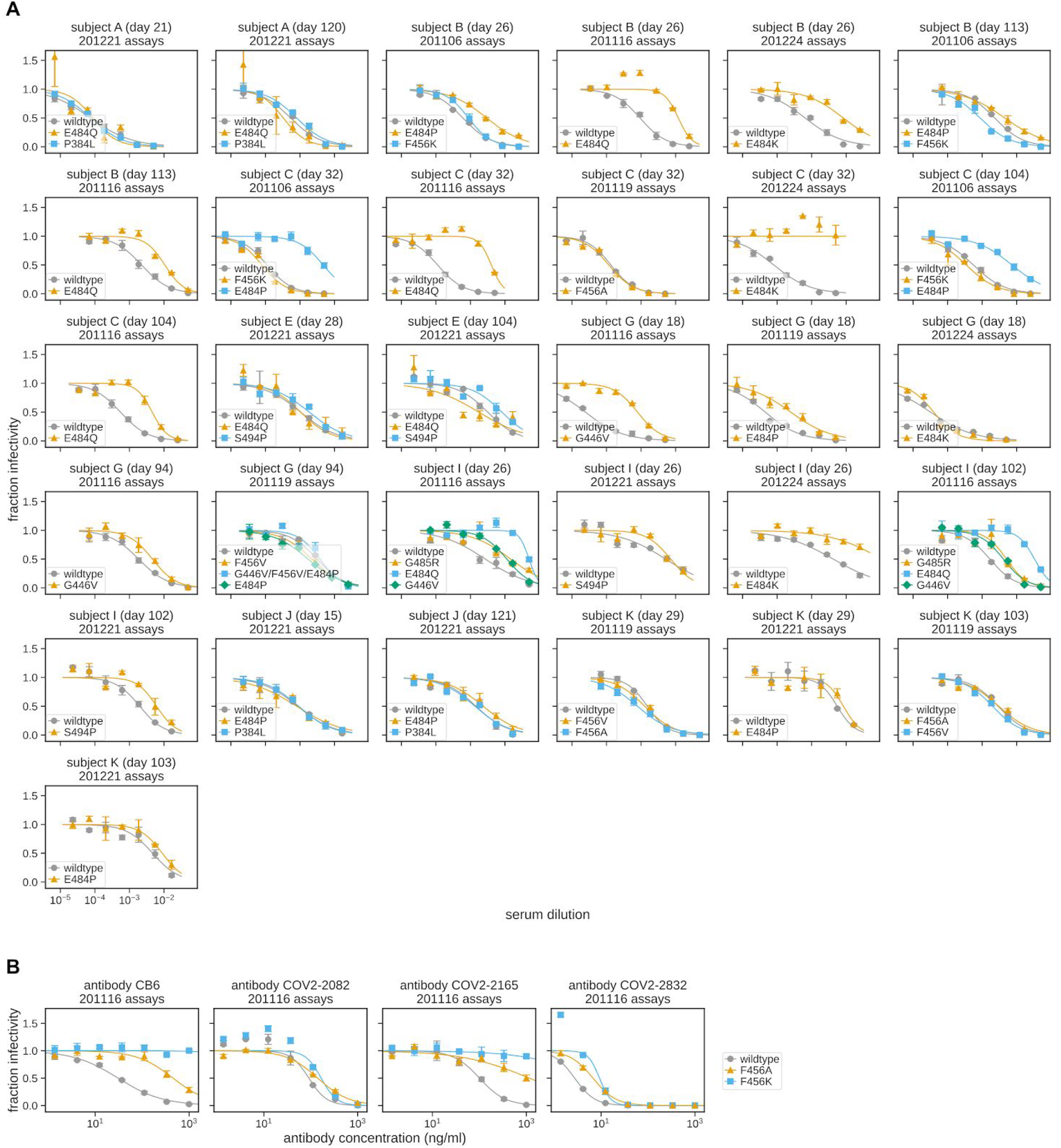
Full curves for all assays testing how RBD mutations affected viral neutralization, related to Figure 4. **(A)** The x-axis gives the serum dilution, and the y-axis gives the fraction of viral infectivity remaining at that dilution. A different plot facet is shown for each serum (labeled by subject and day of collection) and assay date. The neutralization curves were fit and plotted using `neutcurve` (https://jbloomlab.github.io/neutcurve/, version 0.5.1) and fitting 2-parameter Hill curves with the baselines fixed at one and zero to calculate IC50s. These IC50s were then used to determine the fold-change values plotted in **Figure 5A-C**, comparing each mutant to the wildtype run on the same assay date. The curves plotted in **Figure 5D** recapitulate data plotted in this panel, but aggregate mutants across several assay dates and show the wildtype curve for just the first assays date. This aggregation across assay dates is well supported since the wildtype was re-run on each assay date and always yielded very similar IC50s for any given serum. **(B)** Neutralization curves for monoclonal antibodies run against mutations to F456. Our previous escape mapping showed that F456A/K mutations escape binding by the anti-SARS-CoV-2 RBD monoclonal antibodies COV2-2165 and CB6 (also known as LY-CoV016), but not by COV2-2082 or COV2-2832 (Greaney et al., 2020; Shi et al., 2020; Starr et al., 2020a; Zost et al., 2020a). The neutralization assays shown here supported this mapping, and demonstrated that mutations at F456 can indeed greatly reduce neutralization by monoclonal antibodies. The numerical IC50s from all curves in both panels are available at https://github.com/jbloomlab/SARS-CoV-2-RBD_MAP_HAARVI_sera/blob/main/experimental_validations/results/mutant_neuts_results/mutants_foldchange_ic50.csv.

### Supplementary Files

**Supplementary Table 1. Patient metadata and neutralization titers of convalescent sera pre- and post-depletion of RBD-binding antibodies, related to Figure 1**. This table contains the sample metadata from the second supplementary file of Crawford, et al. (2020) as well as the neutralization titers measured in the present study. Each serum is listed twice (on two rows), one with the pre-depletion serological measurements, and one with the post-depletion serological measurements. The table is available online at https://github.com/jbloomlab/SARS-CoV-2-RBD_MAP_HAARVI_sera/blob/main/experimental_validations/results/rbd_absorptions/rbd_depletion_foldchange_ic50.csv

**Supplementary Table 2.** Percentage of RBD mutant library that fell into FACS “escape gate” for each serum during the mapping, related to Figure 2. Serum is a unique identifier for each serum mapped, PID is the patient ID from (Crawford et al., 2020a), subject is the simpler patient identifier used for patients in the current study, the selection serum dilution indicates the reciprocal dilution at which each selection was performed (i.e., 500 is a 1:500 dilution of serum) and the 4 rightmost columns indicate the percentage of each population of cells that fell into the antibody-escape selection gate for the duplicate mutant libraries (lib1 and lib2) and for cells expressing unmutated RBD and incubated with the same dilution of serum as the mutant libraries (WT 1x) or 10-fold less serum (WT 0.1x). There are no corresponding raw FACSDiva gating plots for expt_36 (subject K (day 29)) in **Figure S2**.

| experiment | serum | PID | subject | days post-symptom onset | selection plasma dilution | percentage in escape gate |  |  |  |
| --- | --- | --- | --- | --- | --- | --- | --- | --- | --- |
|  |  |  |  |  |  | lib1 | lib2 | WT 1x | WT 0.1x |
| expt_34 | 23_d21 | 23 | A | 21 | 1250 | 2.6 | 1.7 | 0.2 | 3.2 |
| expt_39 | 23_d45 | 23 | A | 45 | 1250 | 3.1 | 2.5 | 0.1 | 10.8 |
| expt_50 | 23_d120 | 23 | A | 120 | 500 | 4.5 | 6.4 | 0.1 | 27.5 |
| expt_41 | 1C_d26 | 1C | B | 26 | 200 | 4.5 | 3.6 | 0.1 | 0.3 |
| expt_51 | 1C_d113 | 1C | B | 113 | 200 | 3.7 | 4.7 | 0 | 0.4 |
| expt_35 | 24C_d32 | 24C | C | 32 | 200 | 6.7 | 6.5 | 0 | 0.2 |
| expt_44 | 24C_d104 | 24C | C | 104 | 200 | 4.6 | 5.5 | 1 | 1.9 |
| expt_30 | 6C_d33 | 6C | D | 33 | 500 | 5.1 | 4.7 | 0.1 | 27.4 |
| expt_42 | 6C_d76 | 6C | D | 76 | 500 | 4.3 | 3.8 | 0.1 | 1.2 |
| expt_38 | 22C_d28 | 22C | E | 28 | 200 | 4.7 | 2.9 | 0 | 0.9 |
| expt_45 | 22C_d104 | 22C | E | 104 | 200 | 5 | 4.4 | 1.8 | 1.7 |
| expt_48 | 25C_d48 | 25C | F | 48 | 200 | 4 | 5 | 0.2 | 37.7 |
| expt_49 | 25C_d115 | 25C | F | 115 | 80 | 4.4 | 5.9 | 0.2 | 30.8 |
| expt_32 | 25_d18 | 25 | G | 18 | 500 | 5.1 | 5.9 | 0.1 | 3.5 |
| expt_33 | 25_d94 | 25 | G | 94 | 200 | 5.2 | 4.2 | 0.3 | 44.8 |
| expt_37 | 12C_d61 | 12C | H | 61 | 160 | 3.3 | 3.1 | 0 | 0.3 |
| expt_40 | 12C_d152 | 12C | H | 152 | 80 | 4 | 5.2 | 0 | 0.7 |
| expt_46 | 23C_d26 | 23C | I | 26 | 80 | 2.7 | 4.6 | 0 | 1.8 |
| expt_47 | 23C_d102 | 23C | I | 102 | 80 | 3 | 5.8 | 0 | 0.5 |
| expt_31 | 13_d15 | 13 | J | 15 | 200 | 4.4 | 5.7 | 0.1 | 71.3 |
| expt_52 | 13_d121 | 13 | J | 121 | 1250 | 4.2 | 4.8 | 0.1 | 7.2 |
| expt_36 | 7C_d29 | 7C | K | 29 | 500 | 6.1 | 6.1 | 0 | 0.7 |
| expt_43 | 7C_d103 | 7C | K | 103 | 200 | 4.9 | 4.9 | 0.5 | 15.5 |

**Supplementary Table 3. Measurements of the effects of mutations on binding by polyclonal human sera, related to Figure 2**. The file gives the “escape fraction” for each mutation, as well as the total escape fraction at each site and the maximum escape fraction for any mutation at the site. The table is available online at https://raw.githubusercontent.com/jbloomlab/SARS-CoV-2-RBD_MAP_HAARVI_sera/main/results/supp_data/human_sera_raw_data.csv.

## Notes

https://github.com/jbloomlab/SARS-CoV-2-RBD_MAP_HAARVI_sera

https://jbloomlab.github.io/SARS-CoV-2-RBD_MAP_HAARVI_sera/

